# Comparative abundance and diversity of populations of the *Pseudomonas syringae* and Soft Rot *Pectobacteriaceae* species complexes throughout the Durance River catchment from its French Alps sources to its delta

**DOI:** 10.1101/2022.09.06.506731

**Authors:** C.E. Morris, C. Lacroix, C. Chandeysson, C. Guilbaud, C. Monteil, S. Piry, Rochelle Newall E., S. Fiorini, F. Van Gijsegem, M.A. Barny, O. Berge

## Abstract

Rivers, creeks, streams are integrators of biological, chemical and physical processes occurring in a catchment linking land cover from the headwaters to the outlet. The dynamics of human and animal pathogens in catchments have been widely studied in a large variety of contexts allowing the optimization of disease risk reduction. In parallel, there is an emerging awareness that crop pathogens might also be disseminated via surface waters especially when they are used for irrigation. However, there are no studies on the extent to which potential plant pathogens are present – nor about their dynamics - along the full course of a catchment. Here we have compared the seasonal dynamics of populations of the *Pseudomonas syringae* (Psy) and the Soft Rot *Pectobacteriaceae* (SRP) species complexes along a 270 km stretch of the Durance River from the upstream alpine reaches to the downstream agricultural production areas at the confluence with the Rhone River at Avignon. Among 168 samples collected at 21 sites in fall, winter, spring and summer of 2016 and 2017, Psy strains were detected at all sampling sites and in 156 of the samples at population densities up to 10^5^ bacteria L^−1^. In contrast, SRP strains were detected in 98 of the samples, mostly from the southern part of the river, at population densities that did not exceed 3 × 10^4^ bacteria L^−1^. Among the biological and chemical parameters that were characterized at each sampling site, temperature was the only factor that explained a significant amount of the variability in population size for both species complexes. Psy densities decreased with increasing temperature whereas SRP densities increased with increasing temperature. River-borne populations of SRP were composed mainly of *Pectobacterium versatile* and *P. aquaticum* that have little known epidemiological importance. Only a few strains of *Pectobacterium* and *Dickeya* species reputed for their epidemiological impact were observed. In contrast, Psy populations at all sites were dominated by a genetic lineage of phylogroup 2 known from other studies for its broad host range and its geographic and habitat ubiquity. Our observations suggest that surveillance of river water for SRP could be leveraged to signal diagnostic and management reactions to avoid disease outbreaks. In contrast, the constant presence of Psy throughout the catchment in absence of regular and widespread disease outbreaks due to this group of bacteria suggests that surveillance should focus on future changes in land use, river water conditions and agronomic practices that could destabilize the mechanisms currently holding Psy outbreaks in check.

## Introduction

Surface waters are vital components of agro-systems. They provide water for irrigation and industrial processing of foods as well as being important for other uses such as for drinking, generation of electricity, recreation and navigation. Lakes and rivers are defining features of landscape topography and they influence the fertility and humidity of soils in their proximity. Flowing surface waters (rivers, creeks, streams) are physical links between agricultural production fields and other land covers both up- and down-stream as they transport particles and various chemicals that enter rivers along their paths from source to sink. Rivers, lakes, streams and ponds harbor microorganisms that are pathogenic to humans and animals as well as those that are pathogenic to plants (James & Joyce, 2004; Zappia et al., 2014; Lamichhane & Bartoli, 2015). The abundant data on human and animal pathogens in river catchments has led to models of the dynamics of the populations of these microorganisms along the course of rivers. These models are used to assess where water poses risks for human health and where and when to optimize interventions to reduce these risks (Rankinen et al., 2016; Whitehead et al., 2016).

With sufficient knowledge bases, similar applications would be possible for plant health including assessing where use of river water for irrigation poses the greatest risk of plant diseases, conceiving interventions to reduce the risk, and orienting surveillance of river water quality to validate the efficiency of those interventions. In the case of the plant pathogen *Ralstonia solanacearum*, for example, fine scale molecular typing was used to locate aquatic sources of strains of phylotype IIB sequevar 1 attacking potatoes but that are also virulent on other solanaceous crops and weeds (Parkinson et al., 2013). The presence of plant pathogens in water might be due to build-up from cultivation of host plants as suspected in the case of *R. solanacearum* (Tomlinson et al., 2009), whereas others might have – in addition to large host ranges - significant saprophytic phases in water such as *Pseudomonas syringae* (Morris et al., 2010). In this latter case, there are likely to be multiple, diffuse sources of these microorganisms in landscapes rather than discrete sources that can be surmised. Therefore, data are needed all along the course of a river to give models the power to infer sources. Finally, as for human and animal pathogens, data are needed on the regular occurrence of plant pathogens in river water to assure that observations are not anecdotal and that the organism is sufficiently frequent to foster the modeling of its dynamics. In this light, the *Pseudomonas syringae* (Psy) and the Soft Rot *Pectobacteriaceae* (SRPs) species complexes stand out in terms of multiple previous reports of their presence in surface waters (Pérombelon & Kelman, 1980; McCarter-Zorner et al., 1984; Harrison et al., 1987; Eayre et al., 1995; Monteil et al., 2013; Morris et al., 2013; Duprey et al., 2019; Pédron et al., 2019).

Here, we have mapped the abundance of two groups of plant pathogenic bacteria in a 270 km stretch of the Durance River, several tributaries and a canal in Southern France. Situated in a Mediterranean fruit and vegetable production region, the Durance River drains over 14000 km^2^ of which 20% is agricultural production (Andrew & Sauquet, 2017). This river has been exploited since the 1100’s for irrigation, milling, navigation, drinking water, mining of sediment, generation of electricity and recreation. This has involved the creation of canals and dams, restructuration of banks and dredging of sediments leading to changes in flow rates (Andrew & Sauquet, 2017). Land use and ground cover in the Durance River catchment are influenced by the topography of the basin with recreation, pastures and nature reserves mostly in the mountainous zone from its source to the Lake Serre Ponçon reservoir (that retains 1.2 billion m^3^ and is the second largest reservoir in Europe). Downstream of the lake crop cultivation and large urban zones dominate the landscape (Andrew & Sauquet, 2017). This river basin is in a typically Mediterranean region. Therefore, it is subjected to the vicissitudes of climate leading to landslides, flooding and droughts that alter the flow and particle content of the river and that complement the seasonal water discharge dynamics that are mostly influenced by snowmelt.

The objective of this work was to compare the abundance and reoccurrence across seasons of two groups of plant pathogenic bacteria – the *Pseudomonas syringae* and the Soft Rot *Pectobacteriaceae*) species complexes - along the stretch of the Durance River from alpine regions to the agricultural production region where the Durance joins the Rhone River at Avignon. Specifically, we determined the variability in the concentrations and diversity of the populations of these two groups of bacteria at 21 sites across the entire catchment over two years during 4 seasons. We targeted these two groups in light of the ensemble of features that are contrasting and/or overlapping for their ecology and strategies of interaction with host plants. These two plant species complexes are classified within different orders of the gamma-Proteobacteria, the Pseudomonadale order for *P. syringae* and the Enterobacterale order for the SRP. They differ in the mechanisms by which they cause disease - with the SRP secreting a large cocktail of plant cell wall degrading enzyme to destroy the plant cell and recover nutrients and *P. syringae*‘s deploying a type III secretion system that injects a battery of effector proteins into plant cells that collectively allow suppression of plant defenses and gain of access to nutrients (Pérombelon & Kelman, 1980; Lindeberg et al., 2012), suggesting that SRP would likely have greater saprophytic potential than Psy. Nevertheless, they both have very large host ranges collectively. Furthermore, Psy and Pectobacteria are the most frequent causes of new disease emergences in crops among bacterial plant pathogens (Morris et al., 2019). *Pseudomonas syringae* is a species complex composed of numerous phylogroups (PG) and clades (Berge et al., 2014) with a few having recognized taxonomic status as species. In studies that quantify its abundance in the environment (Morris et al., 2010; Monteil et al., 2014; Pietsch et al., 2017; Stopelli et al., 2017) members of this complex are identified based on phylogenetic affiliation according to a partial nucleotide sequence of the citrate synthase housekeeping gene (*cts*). Sequence analysis based on this portion of the *cts* gene allows strain identification and placement in the context of the phylogeny that accounts for the broadest scope of genetic diversity of this group (Berge et al., 2014). Strains in the *P. syringae* group are present in fresh waters and have been isolated previously from sources and tributaries of the Durance River (Morris et al., 2010). However, their abundance along the full course of the Durance River and across seasons has not been assessed. Species of the SRP complex can be quantitatively isolated from environmental sources on a medium that reveals their capacity to degrade pectin (Ben Moussa et al., 2022) and they can be identified based on phylogenetic affiliation according to partial sequences of the housekeeping gene glyceraldehyde-3-phosphate dehydrogenase A (*gapA*) (Cigna et al., 2017). The occurrence of species representing the SRPs throughout the Durance River has been reported recently but not quantitatively (Ben Moussa et al., 2022). Nevertheless, this first report suggests that, despite the capacity of SRPs to proliferate as a saprophyte on decaying plant material making it likely for them to be ubiquitous in rivers (Pérombelon & Kelman, 1980; Jorge & Harrison, 1986; Harrison et al., 1987; Potrykus et al., 2015), the SRPs seem to be markedly different in their population dynamics in river water compared to the ubiquitous *P. syringae* group. Here, we have compared the spatial and temporal dynamics of the populations of these two groups of bacteria to identify the environmental factors and adaptive features that could distinguish them in terms of their capacity to establish reservoirs in river water and especially in rivers used for irrigation of crops.

## Methods

### Sample collection and handling

Water was collected from 21 sites representing the three hydrological sections of the Durance catchment (Kuentz, 2013), of which 8 were along the main course of the river, 11 were from 9 different tributaries that flow into the main river, and 2 were from a major man-made canal (Tab. 1). This canal is a managed distributary of the Durance River and its floodway inlet is located in Mallemort, France (43.73267° N, 5.18599° E). Water was collected at each site at 8 dates to represent four seasons across two years. Sampling campaigns were conducted in 2016 on 1-17 Feb., 13-19 May, 24-28 Aug. and 18-21 Nov; and in 2017 on 3-8 Feb., 4-6 June, 21-25 Aug. and 8-13 Nov. These dates will be referred to, respectively, as Winter-16, Spring-16, Summer-16, Fall-16, Winter-17, Spring-17, Summer-17 and Fall-17.

**Table 1.** Sampling sites in the Durance River catchment.

| Site code<sup>a</sup> | Basin<sup>b</sup> | Latitude °N | Longitude °E | Altitude (m) | Distance from main river (km): |  | Description |
| --- | --- | --- | --- | --- | --- | --- | --- |
|  |  |  |  |  | upstream from confluence | downstream from divergence |  |
| C01a | Lower | 43.756378 | 5.150282 | 106 | - | 5 | Carpentras Canal, a managed distributary of Durance River, near Logis Neuf village |
| C01b | Lower | 43.820640 | 5.082083 | 98 | - | 15 | Carpentras Canal, a managed distributary of Durance River, near Les Taillades village |
| R01 | Upper | 45.024096 | 6.564294 | 1813 | 0 | - | Main course of the river, historically called Clarée River, at this location named “Pont de la Souchère” |
| R02 | Upper | 44.924983 | 6.67987 | 1363 | 0 | - | Main course of the river, historically called Clarée River, at this location named “Pont des amoureux” |
| R03 | Upper | 44.704668 | 6.60111 | 907 | 0 | - | Durance River near St. Crépin village |
| R04 | Upper | 44.550722 | 6.484659 | 790 | 0 | - | Durance River at Embrun city, the entrance to Serre Ponçon Lake |
| R05 | Middle | 44.475576 | 6.112166 | 620 | 0 | - | Durance River at “Archidiacre” site, not far downstream from the Serre-Ponçon Dam. |
| R06 | Middle | 44.212041 | 5.939179 | 459 | 0 | - | Durance River, at « Plan de la Baume » village near Sisteron |
| R07 | Middle | 43.804113 | 5.825458 | 291 | 0 | - | Durance River at Manosque |
| R08 | Lower | 43.667174 | 5.490215 | 188 | 0 | - | Durance River near Pertuis |
| T01 | Upper | 44.924603 | 6.679891 | 1363 | at confluence | - | A tributary of the main river, historically called Durance River, at this location named “Pont des amoureux” and upstream of this confluence |
| T02 | Upper | 45.015400 | 6.459957 | 1659 | 20 | - | Guisane River, a tributary of the Durance river, not far from the “Col du Lautaret” |
| T03 | Upper | 44.877228 | 6.47717 | 1294 | 12 | - | Ailefroide Stream near Pelvoux village, tributary of the Gyr river |
| T04 | Upper | 44.681169 | 6.696794 | 1066 | 9 | - | Guil Stream, a tributary of Durance River, site near the place named “La Maison du Roi”, |
| T05a | Upper | 44.536205 | 6.703007 | 2090 | 55 | - | Riou Mounal Creek, tributary of Ubaye River, site near the “Col de Vars” |
| T05b | Upper | 44.514918 | 6.75597 | 1443 | 50 | - | Ubaye River, a tributary of Durance River, site near Saint-Paul-Sur-Ubaye village |
| T05c | Upper | 44.397539 | 6.480858 | 968 | 11 | - | Ubaye River, a tributary of Durance River, site near Le Martinet village, just upstream of its entrance into Serre Ponçon Lake |
| T06 | Middle | 44.201151 | 5.928711 | 459 | 1 | - | Buëch River, a tributary of Durance river, site near Sisteron |
| T07 | Middle | 44.042293 | 6.040222 | 438 | 4 | - | Bléone River, tributary of Durance River, upstream of a small dam |
| T08 | Middel | 43.728389 | 5.816048 | 274 | 6 | - | Verdon River, tributary of Durance River, site at Vinon-sur Verdon |
| T09 | Lower | 43.884328 | 4.892864 | 39 | 5 | - | Grand Anguillon River, tributary of Durance River, site at Noves where the river becomes a canal |

<sup>a</sup>The codes for the sites indicate where water was collected from the main course of the river (R), a tributary flowing into the main course of the river (T) or a canal used to distribute water from the main course of the river for agricultural and other uses (C).

Surface water was collected at several meters distance from the banks at each site with a 12-L bucket attached to a rope. For all sampling dates, each site was represented by a single bulk sample, resulting in 168 water samples (hereafter referred to as “main experiment”). In Fall-17, triplicate samples at three different times in the day were collected at sites R02 and R08 to assess representativeness of the bulk samples (hereafter referred to as “variability experiment”). The bucket was dedicated to sampling river water, never in contact with high level of bacterial populations. It was rinsed twice with water from the sampled site before collecting the sample. In this way, water from previous sampling sites was washed away. About 1.5 L of water was collected into sterile plastic bottles from the bucket. With the water remaining in the bucket, temperature and electrical conductivity were measured using a Multi Probe System (YSI 556 MPS, YSI, Yellow Springs, USA) and water turbidity was measured using a EUTECH Instruments turbidity meter (TN100, Paisley, Scotland). Samples were maintained in a cooler (ca. 15°C) for no more than 24 h until further processing for chemical and microbiological analyses. To prepare samples for microbiological analyses, 500 mL were filtered across 0.2 µm porosity cellulose acetate filters (Sartorius, 11107-47-ACN, Goettingen, Germany). The bacteria retained on the filter were suspended in 1 mL of sterile distilled water. This suspension, concentrated by a factor of 500 compared to the original sample, was immediately used for subsequent bacterial isolation and quantification. The filtrate was collected for nutrient and dissolved organic carbon analysis as detailed elsewhere (Ben Moussa et al., 2022). Methods for determination of the concentration of DOC, nitrates, nitrites, ammonium, ortho-phosphates, and total dissolved nitrogen and phosphorus were as previously described (Ben Moussa et al., 2022). Briefly, acidified (85% H3PO4), filtered river-water samples (0.2 µm pore diameter) were used to determine the dissolved organic carbon (DOC) concentration with a total organic carbon analyzer (Shimadzu, Kyoto, Japan). The concentration of nitrates, nitrites, ammonium, ortho-phosphates, and total dissolved nitrogen and phosphorus was determined by colorimetry in the laboratory with a segmented continuous flow analyzer (AA3; Seal Analytical, King’s Lynn, UK). The samples (15 ml) were filtered in situ on 0.2 ml for dissolved nutrients and on 50 ml for total nitrogen and phosphorus and frozen (−20° C) before analysis.

### Quantification of total culturable bacteria

The concentrated suspension was dilution-plated on 10% tryptic soy agar as previously described (Morris et al., 2010) and using the same number of replicates and volumes (2-3 technical replicates of 100 µl for each dilution) for quantification of Psy described below. Plates were incubated at ambient temperature (18 to 25°C) for 2 to 4 days. Colonies were counted regularly during the incubation period up to 4 days.

### Isolation and quantification of Soft Rot *Pectobacteriaceae*

The bacterial suspensions were serially diluted in water and plated on crystal violet pectate (CVP) medium plates, a semi-selective medium containing pectin that is widely used for the isolation of pectinolytic *Pectobacterium* and *Dickeya* (Hélias et al., 2012; Faye et al., 2018). Plates were incubated at 28°C for 2 days and the number of colonies forming deep pits in the CVP medium typical of *Pectobacteriaceae* were recorded. For each treated sample, up to 30 pit-forming colonies were purified on CVP medium and further streaked on LB medium for conservation. Qualitative description of the purified strains has been published recently (Ben Moussa et al., 2022). In the present paper, we evaluated the quantity of recovered SRP by counting the deep pits formed on plates and analyzed these data with regard to other variables measured in the course of this study.

### Isolation and quantification of bacteria in the *P. syringae* complex

The concentrated suspension was dilution-plated as previously described (Morris et al., 2010) on King’s medium B supplemented with cephalexin, boric acid and cycloheximide (referred to as KBC medium). Two to three replicates of each dilution were plated to assure that when possible at least 30 colonies suspected to be *P. syringae* (“putative” *P. syringae* based on colony traits) could be isolated for each site at each date. After 3 to 7 days incubation of KBC plates at room temperature (∼20-25°C) the numbers of putative *P. syringae* colonies were recorded. Based on our previous work with the diversity of the *P. syringae* complex (Berge et al., 2014), few phenotypic traits are reliable for screening colonies to hone in specifically on *P. syringae*. Non-putative colonies were eliminated according to pigmentation, pin-point colony size and ornate or crusty colony appearance. From the remaining, 30 or more putative colonies (or all putative colonies if there were fewer than 30) were randomly selected for each sample and streaked onto a plate of King’s medium B (King et al., 1954) to increase the number of bacterial cells per colony for further characterization but not to purify strains. Each isolate was then introduced into the well of 96-well plate (i.e. initial plates) previously loaded with 150 µl of demineralized water kept at 4°C until being identified on the basis of high-throughput MiSeq sequencing of a fragment of the *cts* (citrate synthase) gene and bioinformatic analysis (described below). The sizes of the populations of *P. syringae* in each water sample were calculated by adjusting the number of putative *P. syringae* colonies per each sample according to the percentage that were identified as *bona fide* members of the *P. syringae* complex according to previously established criteria (Berge et al., 2014) through the MiSeq sequencing approach (cf. Supp. Tab. 1 and 3.).

### Preparation of the material for MiSeq sequencing including, putative *P. syringae* isolates, replicates and controls

Among the 5436 isolates introduced into the well of 96-well plates (i.e. initial plates) a total of 537 randomly selected isolates were introduced twice, i.e. into the well of two different 96-well PCR plates, which were further analyzed as replicates to assess the reproducibility of bacterial identification through MiSeq sequencing approach. In addition, each of the initial 96-well PCR plate contained at least one replicate of five different types of controls. Pure colonies of a known *P. syringae* (CC94, phylogroup 02 [PG02]) and *Pseudomonas tolaasi* (CFBP2068) strains were separately introduced into 80 wells of the initial plates (i.e. 160 wells total). Respectively, these two types of positive controls were used to assess the efficiency and repeatability of *P. syringae* identification, and the level of biological or sequencing/bioinformatics contamination across wells. Three types of negative controls were also included in the analysis. A total of 80 and 414 wells were filled only with ultrapure water or PCR mix, respectively (see below), while 142 wells were left empty during the *cts* PCR amplification to check for background contamination from the PCR plate and among wells cross-contaminations (cf. “Amplicon production, preparation of MiSeq libraries and sequencing” paragraph below).

### Amplicon production, preparation of MiSeq libraries and sequencing

The resulting 6769 wells were subjected to PCR amplification targeting a 388 bp fragment (primers excluded) of the *cts* (citrate synthase) gene. PCR was performed on 20 groups of four plates. PCR was performed using a different specifically designed forward primers for each of the 20 groups of four plates (cf. Supp. Tab. 4), and a single common reverse primer (cf. Supp. Tab. 4). Each forward and reverse primers was composed of the binding site for *cts* gene amplification, and of an adapter used in the further steps of MiSeq library preparation for adding the Illumina indexes. In addition, each forward primer included a different 6 nucleotide tag in order to be able to assign output sequences to each initial bacterial isolate during the bioinformatics analyses. The PCR mix was composed of 3.4 µl of 5X Q5 reaction buffer (New England Biolabs), 0.14 µl of 25 mM dNTPs (Promega), 0.68 µl of 10µM of each of the forward and reverse primer, 0.17 µl of Q5 hot start HF DNA polymerase (New England Biolabs), and was adjusted to a final volume of 15 µl with ultrapure water, to which 2µl of the initial material (i.e. isolates, replicates or controls) was added. After a 30 min 98°C activation period of the Taq polymerase, DNA fragments were amplified following 30 cycles of denaturation (10 s at 98°C), annealing (30 s at 55°C) and extension (30 s at 72°C). After a final extension time of 2 min at 72°C, DNA amplicons were stored at –20°C until use. After PCR amplification, four groups of 20 PCR plates were pooled each into a single plate. To this end, 5 µl of the identical well of each of 20 plates were mixed together. The four resulting plates (i.e. pooled plates) were sent to GeT-PlaGe core facility (INRAE, Toulouse, France) where the final MiSeq libraries were prepared. The libraries were run using an Illumina MiSeq pair-end 2*250 pb sequencing technology.

### Bio-informatic processing of raw sequences and identification of Amplicon Sequence Variants

Raw sequences were processed in R (version 4.0.3) (R_Core_Team, 2020) with the FASTA program version 36.3.8h (https://fasta.bioch.virginia.edu/fasta_www2/fasta_list2.shtml), and the packages ShortRead, DADA2 and ggplot2 (Morgan et al., 2009; Callahan et al., 2016; Wickham, 2016). The amplicon and MiSeq library preparation strategy resulted in both forward and reverse reads being present in the R1 and R2 files associated with each well of the pooled plates. Reads pairs were labelled as forward and reverse complement based on the comparison of their sequence with the one of a reference *P.syringae* strain (CVB0016, phylogroup 02 [PG02]; Supp. Tab. 5). Then, read pairs with e-value >= 10^−40^ were removed. Paired reads were sorted such as to include forward and reverse complement reads in final R1 and R2 fastq files, respectively. All reads were demultiplexed using the tags included in the forward primers (cf. Supp. Tab. 4) in order to separate and assign raw sequences to each well of the initial plates. Only sequences that contained exactly matching tags were kept. Reads that were too short, relatively to the required length for merging R1 and R2 reads with an adequate overlap (i.e. 25), were removed. Then, tags and *cts* primer sequences were removed. The quality of sorted and demultiplexed reads was checked and plotted. Reads were not trimmed as the observed error rates were similar to the estimated ones, and as the expected overlap length between the paired forward and reverse reads was relatively short (i.e. 25). Reads that included at least one unidentified nucleotide or which sequence matched the phiX genome were discarded. Then, the 183 Amplicon Sequence Variants (ASVs) were inferred and paired reads were merged. <u>The sequence table that</u> <u>includes the number of copies of paired reads (=ASV) in each well was constructed</u> and chimeras were removed.

Amplicon sequence variants (ASVs) were identified through a blastn+ with the sequence of 910 reference, mostly *P. syringae*, strains (cf. Supp. Tab. 5) using the FROGS Affiliation OTU (Escudié et al., 2018) available on the Genotoul-Sigenae Galaxy server (https://vm-galaxy-prod.toulouse.inra.fr/). The ASVs which percentage identity with the closest reference strains was lower than 98.2% were removed. This value corresponds to the similarity threshold determined previously for accurate clade affiliations within the *P. syringae* species complex (Berge et al., 2014).

### Verification of controls, filters of ASV, and analysis of replicates

A sequence of the expected strain (either *P. syringae* CC94 or *P. tolaasi* CFBP 2068) was identified in 156 out of the 160 wells corresponding to positive controls. No sequence was detected in 3 positive control wells. Some ASVs were identified non-expectedly in one positive control well, and 54 negative control wells, with copies number ranging from 1 to 389. Therefore, a conservative approach was taken whereby each ASV was considered as positively detected in the wells of the initial plates, and thus assigned to the corresponding 5436 isolates, if the number of each pair reads (=ASV) in each well was higher than 400. This resulted in the final identification of 291 *P. syringae* ASV (Supp. Tab 2), which copy number ranged from 400 to 11055 in each initial well where it was detected (hereafter referred as haplotypes). Out of the 537 isolates that were included as duplicates in the initial plates, 392 (73%) yielded in similar results. Specifically, no *P. syringae* haplotype was detected for each of the duplicates of 154 isolates, and a sequence identified as being a *bona fide* member of the *P. syringae* species complex was detected for the duplicates of 238 isolates. The remaining 145 isolates corresponded to cases where a *P. syringae* haplotype was detected for one duplicate but not in the other. Hence, the MiSeq isolate identification approach described here might have led to an under-estimation of the number of *P. syringae* colonies in water samples. However, based on our experience in validating identification of thousands of isolates over the past decades, we have not encountered a case of attribution of identity as *P. syringae* to strains that are clearly outside this species complex. Therefore, we are confident that the likelihood of false positives is rare enough to not be a concern for this work.

### Statistical analyses

The representativeness of single bulk water samples was determined through the analysis of the variability in total Psy and SRP population sizes based on triplicates of water samples taken at three different times of the same day in Fall 2017 at two sites (R02 and R08, Supp. Tab. 3). The analysis was conducted for each site in R (version 4.1.1) (R_Core_Team, 2020) with a linear regression using the log_10_ transformed total number of Psy and SRP colonies per L of water as response variable. The assumption of normality was verified using a Shapiro test (p > 0.62 for site R02 and p = 0.67 for site R08 for Psy; and p > 0.36 for site R02 and p = 0.49 for site R08 for SRP). The variability in population size for both organisms across sampling times within a day (Supp. Fig. 2) was non-significant for both sites (p=0.32 for site R02, p = 0.2 for site R08 for Psy; and p=0.19 for site R02, p = 0.32 for site R08 for SRP).

Statistical analyses of total *P. syringae* population size across the Durance catchment (cf. Supp. Tab. 1) and of associated water chemical characteristics (cf. Supp. Tab. 6) were conducted with the Statistica 10 package (StatSoft www.statsoft.fr, accessed 27 Aug 2019). This included calculation of the correlations among water variables according to Spearman’s Rank correlation and estimation of the contribution of water variables to variability in bacterial concentration (via Principle Component Analysis leading to the construction of composite Principle Component Factors). Statistica was also used to calculate parameters of regressions of the observed values of bacterial population sizes at each site and date (expressed as log_10_ bacteria L^−1^) against single water variables or composite Principle Component factors. Significant effects were reported if p-values were < 0.05.

## Results

### Populations of *P. syringae* and SRP species complexes are present throughout the Durance River catchment but differ in size and frequency of occurrence

Bacterial population sizes were evaluated for 21 sites throughout the Durance River catchment (Tab. 1, Fig. 1). Strains in the *P. syringae* complex (referred to collectively from here on as Psy) were detected at all sampling sites and almost all dates throughout the catchment at population densities up to 10^5^ bacteria L^−1^ (Fig. 1). Population densities of this bacterial group were below the detection threshold (10 - 40 bacteria L^−1^) in only 12 (7%) of the 168 water samples analyzed in this study. In contrast, members of the SRP species complex (referred to collectively from here on as SRP) were less frequently detected than Psy and were most often detected at the sites in the southern-most end of the catchment but rarely in the northernmost reaches of the catchment. SRP population densities were under the detection threshold in 70 (42%) of the samples. SRP and Psy co-occurred in 87 (52%) of the samples. When there was co-occurrence of SRP and Psy, SRP population densities were equal to or exceeded those of Psy in 25% (22) of those samples by up to about one order of magnitude; Psy population densities exceeded those of SRP in 75% (65) of those samples by up to nearly four orders of magnitude (Supp. Fig. 1).

**Figure 1.**
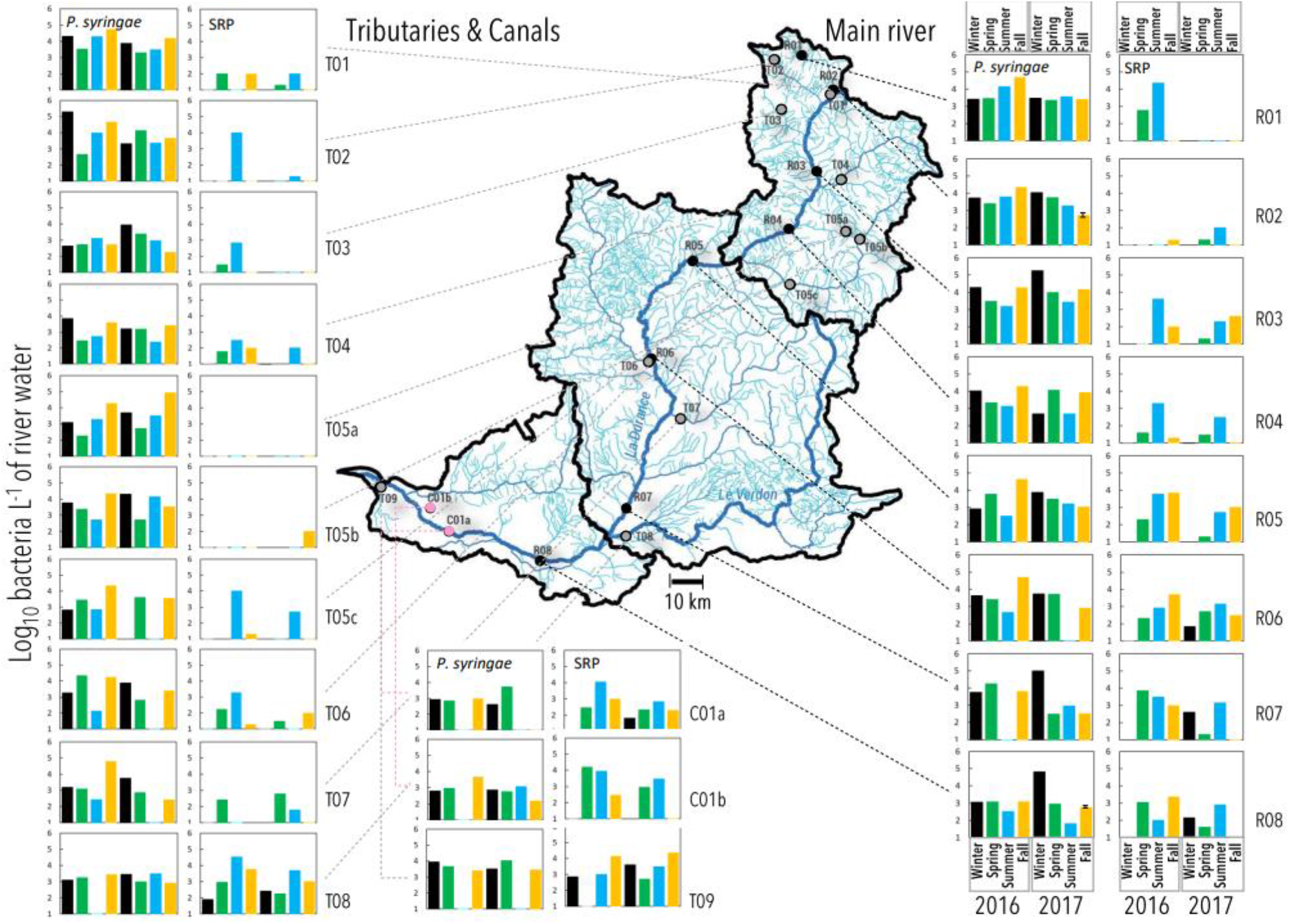
Population densities (log_10_ bacteria L^−1^) of *Pseudomonas syringae* and Soft Rot *Pectobacteriaceae* (SRP) in water in the Durance River basin at eight sites in the main river (R), eleven sites in tributaries (T) and two sites in irrigation canals (C) in four seasons (Winter in black, Spring in green, Summer in blue and Fall in gold) in each of 2016 and 2017. All values for population density are based on culturable bacteria isolated from single samples at each date and site except for values for sites R02 and R08 in 2017 that were means of triplicate samples. Error bars for those values represent the standard error. The map portrays the sampling sites along the full expanse of the Durance River basin from its most northern reaches in the Hautes Alpes department southward through the departments of Alpes d’Haute Provence and Vaucluse. The black contours of the map represent the three hydrological sections of the river basin (Kuentz, 2013). The Durance and Verdon rivers are labeled on the map.

Among the different sampling dates and sites, total culturable populations ranged from 10^5^ to 5 × 10^7^ bacteria L^−1^ (Supp. Tab. 1). Although there was an overall positive trend in the correlation between the densities of total culturable bacterial populations and those of Psy or SRP, the statistical significance (at the 5% level) of the correlation depended on the geographic location according to the three basins of the catchment (delimited in Fig. 1). In the upper, northernmost basin, densities of Psy and SRP were each significantly correlated with total population densities (R = 0.275 and 0.222 for Psy and SRP respectively; p = 0.009 and 0.038, respectively). In the southernmost, lower basin the population densities of neither bacterial group were significantly correlated with total population density (R = 0.187 and 0.052 for Psy and SRP respectively; p = 0.304 and 0.779, respectively). The middle basin differentiated Psy from SRP where total population densities were significantly correlated with Psy densities (R = 0.340, p = 0.018) but not with SRP densities (R = 0.245, p = 0.094). These results suggest that the factors that influence the densities of Psy and SRP populations are likely to be somewhat different from each other and not completely correlated with the factors influencing the abundance of total culturable bacteria.

### Among variables describing the physical-chemical conditions of river water, temperature has the greatest predictive power for population sizes of Psy and SRP with inverse effects on these two species complexes

Seven variables describing the physical-chemical characteristics of the water at each sampling time and according to the geographical context of the site (altitude, longitude and latitude) were measured. Temperature, conductivity and dissolved organic carbon (DOC) concentration were measured in 2016 and 2017 and, in addition, concentrations of PO ^3-^, NH_4_ ^+^, NO ^−^ and NO ^−^ were determined in 2017. The ranges of values for these variables are presented in Fig. 2 and are indicative of an alpine catchment with increasing influence of human activities and agriculture as altitude decreases. There were few correlations among variables for water conditions (Tab. 2, for the ensemble of water variables assessed in 2017). Therefore, we could not eliminate any variables with obvious redundancy to simplify the analysis of their contribution to bacterial population size. Nevertheless, to assess the influence of the ensemble of the physical-chemical properties on bacterial population size, we used Principle Component Analysis (PCA) to construct composite factors that accounted for the importance of each of the seven individual physical-chemical variables determined in 2017 for the overall variability of water conditions (Tab. 3).

**Figure 2.**
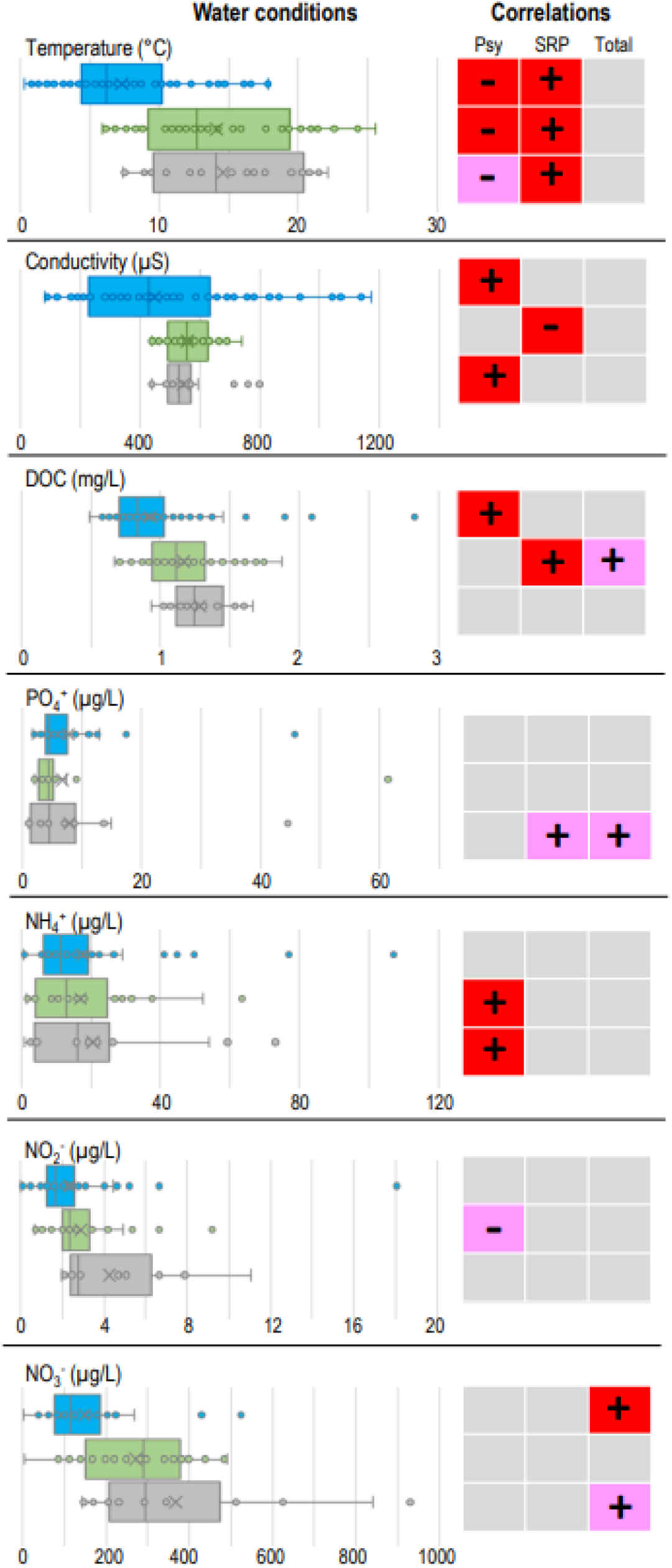
Correlation of *Pseudomonas syringae* (Psy), Soft Rot *Pectobacteriaceae* (SRP) and total mesophilic (Total) bacterial population densities with water conditions in the three basins of the Durance River catchment. The left-hand panel indicates the water conditions (box plots including a presentation of all data values) in the three basins (as depicted in Fig. 1) (upper in blue, middle in green and lower in grey). The box plots represent the median (black line), the mean (X), the 25%-75% quartile (the box) and the minimum and maximum values under the assumption of a normal distribution of data that would exclude outliers (whiskers). The right hand panel indicates whether the values of the Spearman Rank correlation between the water conditions and each of the bacterial population densities were positive (+) or negative (-), and if they were significant according to p < 0.05 (red background) or 0.05 > p < 0.10 (pink background). Grey backgrounds indicate that p > 0.10 for this statistical test.

**Table 2.**
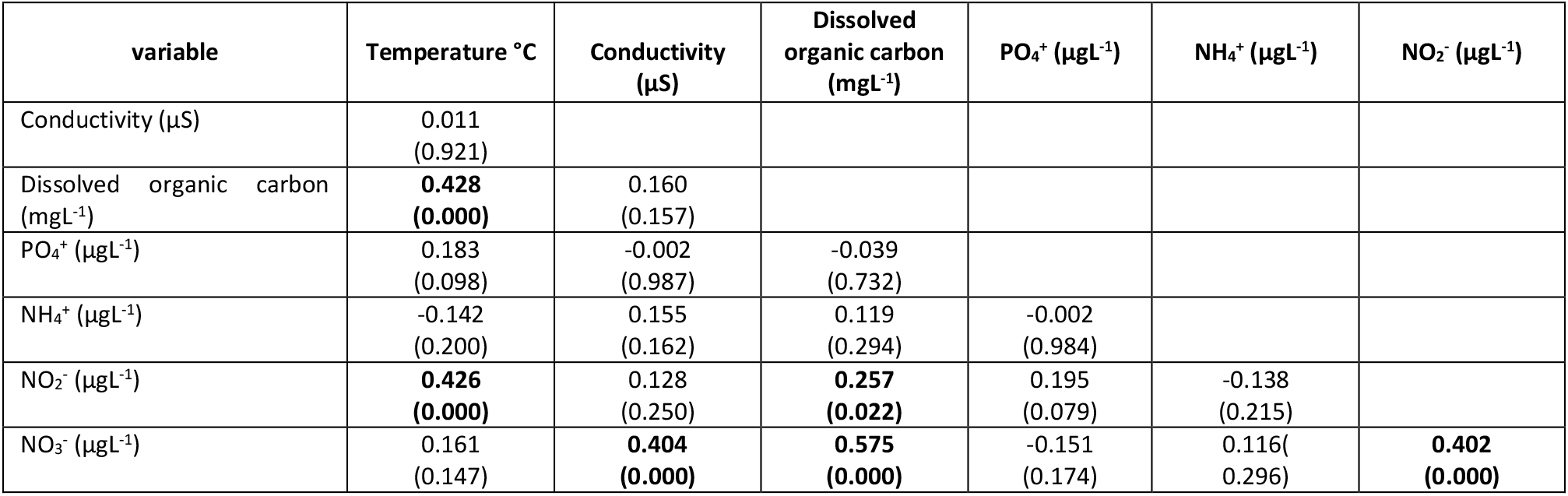
Spearman rank correlations (and associated p-values) among variables describing physical-chemical conditions of water collected at 8 dates at each of 21 sites throughout the Durance River catchment. Significant correlations (p < 0.05) are in bold face.

| variable | Temperature °C | Conductivity ($\mu S$) | Dissolved organic carbon ($mgL^{-1}$) | $PO_4^+$ ($\mu gL^{-1}$) | $NH_4^+$ ($\mu gL^{-1}$) | $NO_2^-$ ($\mu gL^{-1}$) |
| --- | --- | --- | --- | --- | --- | --- |
| Conductivity ($\mu S$) | 0.011<br>(0.921) | | | | | |
| Dissolved organic carbon ($mgL^{-1}$) | **0.428**<br>(**0.000**) | 0.160<br>(0.157) | | | | |
| $PO_4^+$ ($\mu gL^{-1}$) | 0.183<br>(0.098) | -0.002<br>(0.987) | -0.039<br>(0.732) | | | |
| $NH_4^+$ ($\mu gL^{-1}$) | -0.142<br>(0.200) | 0.155<br>(0.162) | 0.119<br>(0.294) | -0.002<br>(0.984) | | |
| $NO_2^-$ ($\mu gL^{-1}$) | **0.426**<br>(**0.000**) | 0.128<br>(0.250) | **0.257**<br>(**0.022**) | 0.195<br>(0.079) | -0.138<br>(0.215) | |
| $NO_3^-$ ($\mu gL^{-1}$) | 0.161<br>(0.147) | **0.404**<br>(**0.000**) | **0.575**<br>(**0.000**) | -0.151<br>(0.174) | 0.116<br>(0.296) | **0.402**<br>(**0.000**) |

**Table 3.**
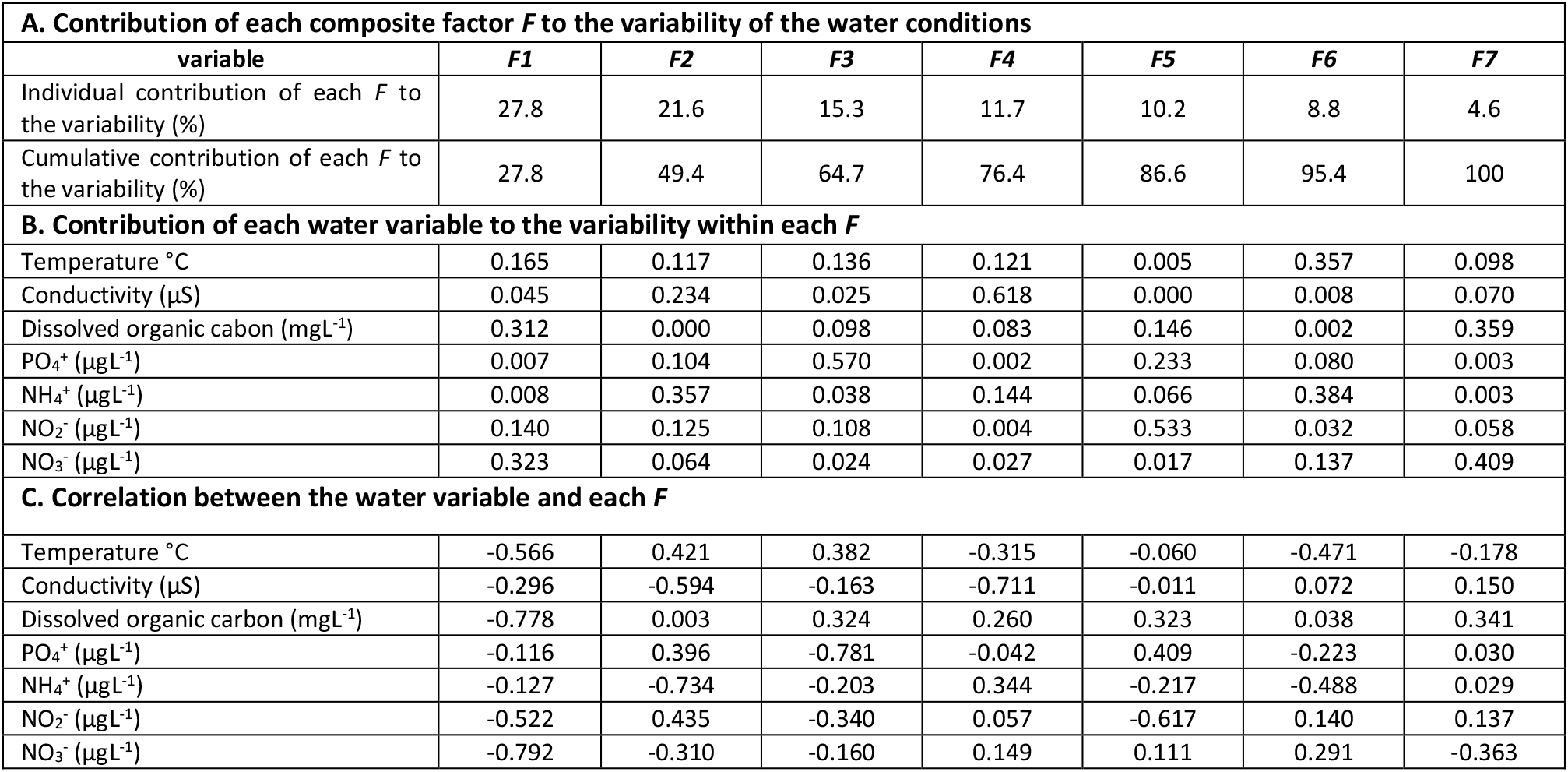
Description of composite factors (*F*) from Principal Component Analysis of seven variables of the physical-chemical conditions of water collected at 8 dates at each of 21 sites throughout the Durance River catchment in terms of (A) the contribution of each *F* to the overall variability of the water conditions, (B) the contribution of each individual variable to the variability within each *F* and (C) the correlation of each water variable with each *F*.

| A. Contribution of each composite factor *F* to the variability of the water conditions |  |  |  |  |  |  |  |
| --- | --- | --- | --- | --- | --- | --- | --- |
| variable | *F1* | *F2* | *F3* | *F4* | *F5* | *F6* | *F7* |
| Individual contribution of each *F* to the variability (%) | 27.8 | 21.6 | 15.3 | 11.7 | 10.2 | 8.8 | 4.6 |
| Cumulative contribution of each *F* to the variability (%) | 27.8 | 49.4 | 64.7 | 76.4 | 86.6 | 95.4 | 100 |
| B. Contribution of each water variable to the variability within each *F* |  |  |  |  |  |  |  |
| Temperature °C | 0.165 | 0.117 | 0.136 | 0.121 | 0.005 | 0.357 | 0.098 |
| Conductivity (µS) | 0.045 | 0.234 | 0.025 | 0.618 | 0.000 | 0.008 | 0.070 |
| Dissolved organic carbon (mgL<sup>-1</sup>) | 0.312 | 0.000 | 0.098 | 0.083 | 0.146 | 0.002 | 0.359 |
| PO<sub>4</sub><sup>3-</sup> (µg L<sup>-1</sup>) | 0.007 | 0.104 | 0.570 | 0.002 | 0.233 | 0.080 | 0.003 |
| NH<sub>4</sub><sup>+</sup> (µg L<sup>-1</sup>) | 0.008 | 0.357 | 0.038 | 0.144 | 0.066 | 0.384 | 0.003 |
| NO<sub>2</sub><sup>-</sup> (µg L<sup>-1</sup>) | 0.140 | 0.125 | 0.108 | 0.004 | 0.533 | 0.032 | 0.058 |
| NO<sub>3</sub><sup>-</sup> (µg L<sup>-1</sup>) | 0.323 | 0.064 | 0.024 | 0.027 | 0.017 | 0.137 | 0.409 |
| C. Correlation between the water variable and each *F* |  |  |  |  |  |  |  |
| Temperature °C | -0.566 | 0.421 | 0.382 | -0.315 | -0.060 | -0.471 | -0.178 |
| Conductivity (µS) | -0.296 | -0.594 | -0.163 | -0.711 | -0.011 | 0.072 | 0.150 |
| Dissolved organic carbon (mg L<sup>-1</sup>) | -0.778 | 0.003 | 0.324 | 0.260 | 0.323 | 0.038 | 0.341 |
| PO<sub>4</sub><sup>3-</sup> (µg L<sup>-1</sup>) | -0.116 | 0.396 | -0.781 | -0.042 | 0.409 | -0.223 | 0.030 |
| NH<sub>4</sub><sup>+</sup> (µg L<sup>-1</sup>) | -0.127 | -0.734 | -0.203 | 0.344 | -0.217 | -0.488 | 0.029 |
| NO<sub>2</sub><sup>-</sup> (µg L<sup>-1</sup>) | -0.522 | 0.435 | -0.340 | 0.057 | -0.617 | 0.140 | 0.137 |
| NO<sub>3</sub><sup>-</sup> (µg L<sup>-1</sup>) | -0.792 | -0.310 | -0.160 | 0.149 | 0.111 | 0.291 | -0.363 |

The PCA led to the construction of seven composite factors (*F*1 – *F*7) for the 79 observations in 2017, each based on a complete set of observations for all water variables. Five factors were needed to explain at least 80% of the overall variability of water conditions (Tab. 3A). Water temperature contributed ca. 10% to 30% of the variability of six of the factors and the other water variables contributed to the same extent of variability for four or fewer of the factors (Tab. 3B). A multiple regression of the population sizes of either Psy, SRP or total mesophilic bacteria against all seven factors was conducted to assure that the potential impact on bacterial population sizes was assessed in light of the infrequent correlation among the water variables (Tab. 2). This revealed a significant contribution of *F*1, *F*2 and *F*4 to the variability of Psy population sizes; a significant contribution of *F*1, *F*3 and *F*6 to the variability of SRP population sizes; and a significant contribution of *F*1 and *F*2 to the variability of total bacterial population sizes throughout the catchment and across seasons in 2017 (Tab. 4). We note that F6 makes a significant contribution to explaining the variability of SRP population sizes although it was outside of the factors that explained at least 80% of the variability of the water conditions. To identify the water variables that contributed the most to the variability of Psy, SRP and total bacterial populations, we ranked the contribution of each water variable for each *F* (Tab 3B) and calculated the cumulative contribution to the variability of each *F* with decreasing rank. For *F*1, *F*3, *F*4 and *F*6, three water variables explained at least 80% of the variability of the factors; and for F2, four water variables explained at least 80% of the variability of each factor (here, we refer to these as the “top explanatory variables”). Among the top explanatory variables, only temperature was common to all the *F* that had significant effects in the regressions (i.e., *F*1, *F*2, *F*3, *F*4 and *F*6). For the *F* that were significant for Psy populations, conductivity and the concentration of NH_4_^+^ were common to two factors (*F*2, *F*4); likewise for SRP the concentration of NO_3_^−^ was common to two factors (*F*1, *F*6). Otherwise, there were no other top explanatory variables that were consistently common to the significant *F* factors.

**Table 4.** Parameters from the multiple regression of population sizes of Psy or SRP (expressed as log10 bacteria L^−1^) vs seven composite factors (*F*) from Principal Component Analysis (*c.f.* Tab. 3). Significant values are in bold face.

| *F* | Dependent variable: log<sub>10</sub> Psy L<sup>-1</sup><br>R = 0.541, R<sup>2</sup> = 0.293<br>p-value<sub>regression</sub> = **0.000** |  | Dependent variable: log<sub>10</sub> SRP L<sup>-1</sup><br>R = 0.628, R<sup>2</sup> = 0.394<br>p-value<sub>regression</sub> = **0.000** |  | Dependent variable: log<sub>10</sub> Total L<sup>-1</sup><br>R = 0.491, R<sup>2</sup> = 0.241<br>p-value<sub>regression</sub> = **0.004** |  |
| --- | --- | --- | --- | --- | --- | --- |
|  | Slope (b) | p-value | Slope (b) | p-value | Slope (b) | p-value |
| 1 | **0.183** | **0.034** | **-0.583** | **0.000** | **-0.125** | **0.000** |
| 2 | **-0.352** | **0.000** | 0.046 | 0.594 | **-0.103** | **0.010** |
| 3 | -0.092 | 0.422 | **0.265** | **0.011** | 0.012 | 0.792 |
| 4 | **0.380** | **0.005** | -0.152 | 0.195 | 0.016 | 0.768 |
| 5 | -0.021 | 0.882 | -0.156 | 0.213 | 0.088 | 0.124 |
| 6 | 0.201 | 0.183 | **-0.293** | **0.031** | 0.008 | 0.892 |
| 7 | 0.214 | 0.304 | -0.338 | 0.072 | 0.036 | 0.673 |

In light of the dominant correlation of water temperature with the size of Psy, SRP and total bacterial populations, we determined to what extent this variable alone could explain the variability in population size. Simple linear regressions of bacterial population size vs temperature revealed that temperature alone significantly explained about 20-40% of the variability of the population sizes of Psy and SRP in 2017 (R^2^ = 0.187, p = 0.000 for Psy; R^2^ = 0.393, p = 0.000 for SRP). When both 2016 and 2017 were considered together, temperature explained about 25-35% of the variability of the sizes of these bacterial populations (R^2^ = 0.249, p = 0.000 for Psy; R^2^ = 0.360, p = 0.000 for SRP). In contrast, temperature alone had no significant explanatory power for the variability of total bacterial population sizes in 2017 (R^2^ = 0.018, p = 0.223) and explained only 5% of the variability of total bacterial populations when 2016 and 2017 sampling campaigns were considered together (R^2^ = 0.050, p = 0.004).

Whereas temperature explained about the same amount of variability of Psy and SRP population sizes, it had inverse effects on population size (Fig. 2). For both Psy and SRP, a change of 10°C was associated with roughly a change in population size by a factor of 10. In the case of Psy, populations increased with decreasing temperature; in the case of SRP, populations decreased with decreasing temperature within the range of temperatures observed in this study. The regression for SRP population sizes predicts that populations would be below the detection level when water temperatures are less than 7°C (Fig. 2). In this study, there were 55 observations where water temperature was colder than 7°C. For these 55 cases, SRP populations were below the detection level for 37 cases whereas Psy populations were detected for all of these cases.

The overriding correlation of temperature with densities of Psy and SRP populations might be due in part to the effect of the wide range of temperatures that are accounted for when data were pooled from sites across the three basins (Fig. 1) spanning altitudes from 39 m to 2090 m (Tab. 1). Pooling data from sampling sites across the three basins might also mask local effects of other water variables that are affected by increasing anthropogenic activities along the land use from the source to the delta of the Durance River catchment. Therefore, we assessed the correlations of bacterial populations with water variables for each of the three basins separately (basin attribution is indicated in Tab. 1). The mean temperature of the water in the upper basin during the sampling campaigns was about 7 °C cooler than that of the middle and lower basins (Fig. 2). Nevertheless, population densities of SRP were positively and significantly (p < 0.05) correlated with temperature in each of the three basins (Fig. 3). Likewise, population densities of Psy were negatively correlated with temperature in each of the three basins; correlations were significant at the 5% level for the upper and middle basins and at the 10% level for the lower basin. Among the top explanatory variables identified above via PCA, NH_4_ ^+^ concentrations were significantly and positively correlated with Psy population densities in the middle and lower basins in spite of the similar concentrations of this compound across the three basins (Fig.2). Although NO_3_ ^−^ concentrations were identified in PCA as one of the top explanatory water variables for SRP densities, there were no significant correlations at the 5% level in any of the three basins. In the PCA, neither conductivity nor DOC were identified as important explanatory factors for the variability of bacterial populations. Nevertheless, conductivity was positively correlated (p < 0.05) with densities of Psy in the upper basin and in the lower basin. These basins were markedly different in the range of conductivity values observed. In the middle basin, where the range of conductivity was similar to that of the lower basin, this variable was negatively correlated (p < 0.05) with SRP densities but had no significant correlation with Psy densities. DOC was positively correlated (p < 0.05) with Psy densities in the upper basin and with SRP densities in the middle basin but not elsewhere.

**Figure 3.**
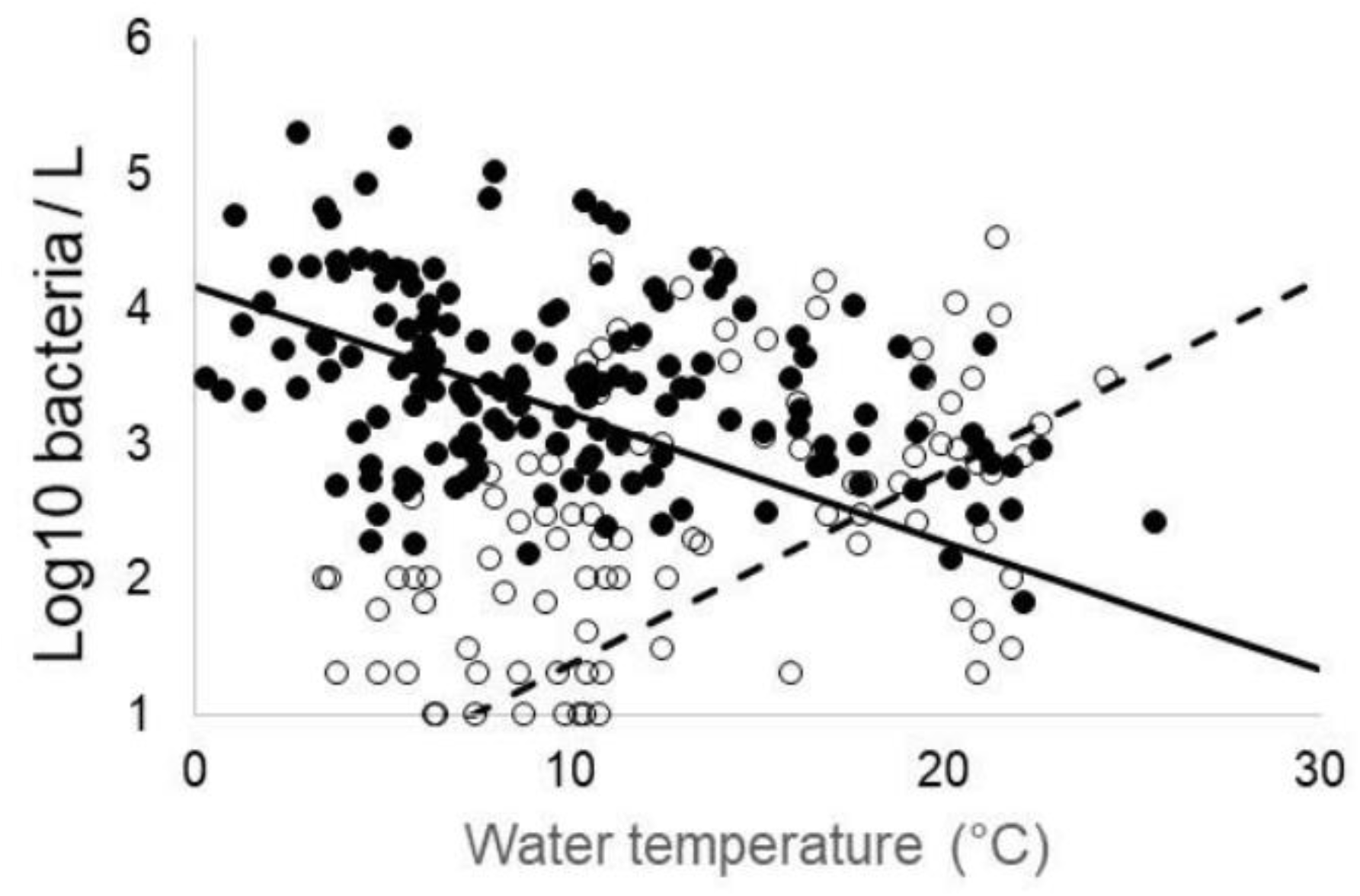
The relationship of bacterial population sizes with water temperature in the Durance River catchment. Water temperature accounted for about 30% of the variability in population size of Psy (solid symbols) (R^2^ = 0.277, p_regression_ = 0.000) and SRP (open symbols) (R^2^ = 0.317, p_regression_ = 0.000) according to linear regressions for data from both 2016 and 2017 combined. The linear regressions are represented by a solid line for Psy (Log_10_ Psy L^−1^ = 4.248 – 0.100 × °C) and a dotted line for SRP (Log_10_ SRP L^−1^ = 0.346 – 0.136 × °C).

The correlations of total population sizes with water conditions were in marked contrast to those for Psy and SRP populations. When assessed according to the individual basins, there were no consistent correlations of total populations with any water variables in a basin with the exception of NO_3_^−^ concentrations (Fig. 3). Total bacterial population densities were positively correlated with NO_3_^−^ concentrations in both the upper and lower basins.

### Populations of *P. syringae* and SRP species complexes in the Durance River catchment are composed of both pandemic and endemic genotypes representing bacterial groups with and without known pathogenic potential

As previously reported (Ben Moussa et al., 2022), the 582 SRP strains isolated from the catchment and identified at species level based on multi-locus sequence typing analysis were composed of *Pectobacterium* (94% of SRP strains from the Durance catchment) and *Dickeya* (6%) species. *Pectobacterium* populations were dominated by species that have no reported epidemiological importance including *P. versatile* (known to be associated with a wide range of plants) and *P. aquaticum* (not known to be pathogenic) constituting 47% and 40%, respectively, of the *Pectobacterium* strains isolated (Ben Moussa et al., 2022). In contrast, important *Pectobacterium* pathogens described on crop such as *P. atrosepticum* or *P. brasiliense* were rarely detected or absent. For *Dickeya* populations (6% of the SRP population), *D. oryzae* (pathogenic mainly on monocots but also on potato) constituted 72% of the *Dickeya* isolates. Among the few remaining *Dickeya* strains, all belong to species of known epidemiological importance including *D. oryzae*, *D. fangzhongdai*, *D. solani*, *D. dianthicola*, and *D. dadantii.* (Ben Moussa et al., 2022).

For Psy, identification was based on phylogroups (PG) and haplotypes within PGs. For these PGs and haplotypes we could then associate them with likely epidemiological behaviors based on previous descriptions. Phylogenetic characterization was conducted for 5436 colonies isolated here that were putative Psy. For these colonies, based on criteria described in material and methods, 2628 could be attributed to known phylogroups of Psy based on comparison with a 388 bp segment of the *cts* gene for 910 strains in the reference data set, and were used to study Psy diversity. The strains that were not attributed to known phylogroups of *P. syringae* might indeed be within the *P. syringae* complex but they were not included in the analyses here because of current taxonomic uncertainties. Strains were identified as belonging to PG01 (9.34 % of all strains), PG02 (45.04 %), PG03 (0.16 %), PG04 (1.30 %), PG07 (13.68 %), PG09 (8.00 %), PG10 (11.60 %), PG12 (0.05 %), PG13 (10.12 %) and PG15 (0.57 %). In contrast to *SRP* species where only *P. versatile* was distributed throughout the catchment and other species were mostly in the southern part of the catchment (Ben Moussa et al., 2022), six PGs of Psy (PG01, 02, 7, 9, 10 and 13) were detected at 19-21 of the 21 sampling sites. The other PGs that each constituted only about 1% or less of the Psy population were found at fewer sites (at 13 sites for PG04 and PG015; two sites for PG03 and PG12).

The 2628 strains of Psy represented 291 different *cts* sequences (referred to here as haplotypes). Nearly half of the haplotypes were endemic in that each was found only at a single sampling site. There were 128 haplotypes with this limited, endemic distribution (Fig 4). Nevertheless, these rare haplotypes only accounted for 5% (154 strains) of the 2628 strains assigned to known PG. Overall, 18 haplotypes accounted for 50% of these strains and each were detected at 15 or more sampling sites. Among these haplotypes, one (referred to here as DD.1) was detected at all 21 sampling sites and represented 10% of all of the strains attributed to known PG in this study. Among individual samples, the fraction of the total population of Psy that was constituted by DD.1 was very consistent and showed a strong positive correlation between the size of the Psy population and that of DD.1 (Spearman Rank correlation coefficient = 0.917, p = 0.000).

**Figure 4.**
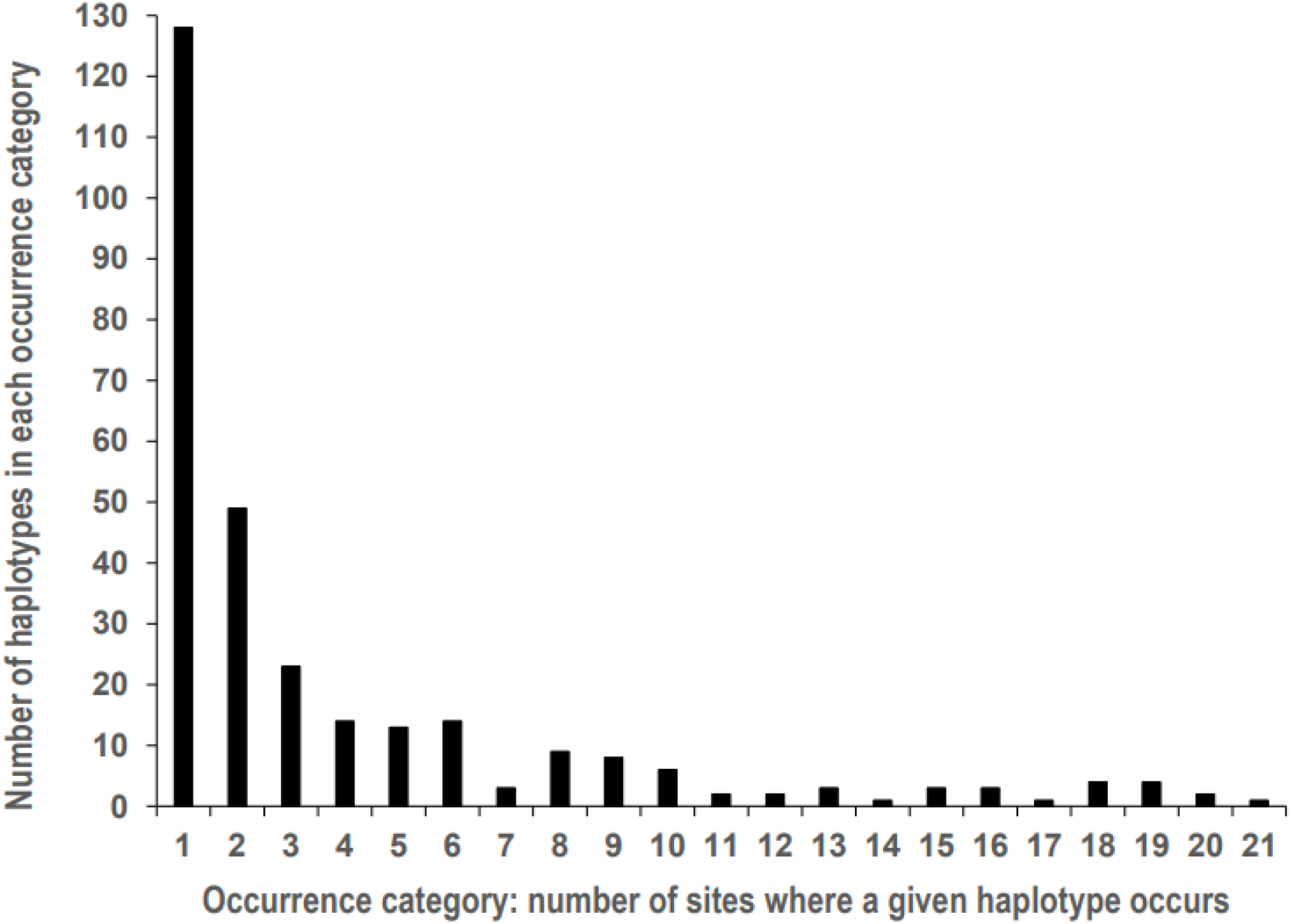
Frequency of occurrence of the 291 haplotypes of *P. syringae* throughout the Durance River basin. Only one haplotype of *P. syringae* (DD.1) was detected at all 21 sampling sites whereas 128 haplotypes were detected at only 1 site during the two years of sampling.

We found that haplotype DD.1 corresponds to a *cts* haplotype of PG02 (in the PG02b clade) that is very widespread when compared to previous reports (Morris et al., 2010; Morris et al., 2019). This *cts* haplotype was the same (100% identity of the 388 bp segment. The *cts* sequence is listed among the amplicon sequence variants in Supp. Tab. 2) as the dominant Psy haplotype found among the 236 strains isolated from river headwaters sampled in the US (Montana and Wyoming), Europe (mostly France and Italy) and New Zealand, representing 39 of the water strains and being the only haplotype found on all three continents and at 11 of the 13 sites sampled (Morris et al., 2010). When compared to reference strains in the study of headwaters by Morris and colleagues (Morris et al., 2010), the haplotype that dominated in headwaters is also the same as the haplotype of 15 reference strains from crops used in that study. These included the type strain of *P. syringae* pv. *syringae* (CFBP1392) isolated from lilac in the UK in 1950 (Gardan et al., 1991), strains 601 and CFBP1906 of *P. syringae* pv. *aptata* isolated from sugar beet in Japan in 1966 (Sarkar & Guttman, 2004) and in France in 1979 (Guillorit-Rondeau et al., 1996), respectively, and strains involved in an epidemic of bacterial blight of cantaloupe that was described to have emerged in France as of 1993 (Morris et al., 2000) (strains CC0001, CC0037, CC0125, CC0354, CC0440, CC0441, CC0457). Additional strains in this haplotype were identified in a subsequent study on host range of Psy (Morris et al., 2019) showing that strains with the same *cts* haplotype as DD.1 were involved in diseases of apricot in France (strain 41A, according to strain names indicated by Morris and colleagues (Morris et al., 2019)), of cantaloupe and squash in New Zealand (CFBP 1788, ICMP 3390, ICMP 7501), and of sugar beets in the Netherlands (CFBP 2471 and CFBP 2507) and Serbia (P004 – P102). By comparing the 388 bp *cts* sequence of haplotype DD.1 to the full GenBank database (BLAST search, https://www.ncbi.nlm.nih.gov/) we also found 100% identity with strains from a freshwater lake in Virginia (strains CLC07, CLC10 (Pietsch et al., 2017)), from freshly fallen snow collected at a high altitude meteorological observatory in Switzerland (JFJ-0007, JFJ-0043 (Stopelli et al., 2017)), from blighted leaves of pea in Japan (H5E3 (Sarkar & Guttman, 2004)) and from home garden philodendrons (several IZB1 and IZB2 strains (Ivanović et al., 2018)) as documented examples that expand the sites and substrates of isolation compared to the information described earlier in this paragraph.

## Discussion

Although there have been previous reports of *P. syringae* and Soft Rot *Pectobacteriaceae* complexes in surface waters (Morris et al., 2010; Toth et al., 2021) here we have made comparisons of their prevalence and abundance across a range of sites representing the diverse environmental conditions across a catchment. This comparison is intended to lead us to identify what typifies each system and what trends are shared. In our effort to make quantitative comparisons of the two groups of bacteria, we faced an initial challenge due to the difference in magnitude of the number of isolates we could collect for each group of bacteria. In a previous study, we verified that differences in abundance were not due to differences in isolation efficiency of the two media used in this work (Pédron et al., 2020). Whereas the hundreds of SRP collected could be characterized as individual strains (sequencing of single or multiple housekeeping genes or full genomes), the thousands of Psy isolates encountered led us to adopt a high-throughput MiSeq sequencing strategy of amplicons of a single housekeeping gene on the basis of Psy-like colonies isolated on KBC growing media. By overcoming this technical challenge, we have shown that for both bacterial species complexes there are genetically diverse populations present throughout the full expanse of the Durance River catchment from near its source - across varying topography, altitude and land use - to the delta where it converges with the Rhone River. Among the SRP, *Pectobacterium* spp. were the most frequently encountered representatives with *P. versatile* being present throughout the catchment (Ben Moussa et al., 2022). Psy populations were dominated by a *cts* haplotype that accounts for 10% of populations at all sites and sampling dates. The structure of Psy populations in the Durance River catchment is similar to that of *Listeria monocytogenes* in surface waters (lakes, rivers, ponds) along the Central California Coast that is dominated overall by a clonal line that constituted 27% of the 1200 strains isolated from these waters (Gorski et al., 2022). However, the specific clonal dominating *L. monocytogenes* populations in waters or other environmental reservoirs differed among the different geographic locations studied (Gorski et al., 2022). In contrast, we observed that Psy populations in distant geographical locations are dominated by the same haplotype that is dominating the Durance River populations overall (Morris et al., 2010), thus illustrating the efficiency of Psy dissemination and the overriding capacity for the DD.1 haplotype to dominate Psy populations.

Populations of Psy were detectable at nearly all sites and all seasons during the two years of sampling whereas SRP populations were frequently below the detection threshold and especially in the upper basin of the catchment where they could not be detected in more than half of the samples. In light of the marked saprophytic capacity of many of the soft rotting bacteria (Pérombelon & Kelman, 1980) and numerous reports of their presence in surface waters (Toth et al., 2021) it could be considered surprising that we did not detect SRP more frequently than Psy. However, our observations suggest that temperature adaptation has a critical role in the ecology of the Psy and SRP species complexes. Our observations also reinforce the idea that Psy is particularly well adapted to freshwater habitats as well as the various other habitats and substrates (plants, precipitation, litter) that are linked via the dissemination of *P. syringae* through the hydrological cycle (Morris et al., 2013). In contrast, SRP are likely to be more dependent on proximity of and seasonality of external plant sources (Toth et al., 2021). In comparison, for *Listeria monocytogenes* that has potential to be a saprophyte as well as a human pathogen (Gorski et al., 2022), the relative importance of anthropogenic *vs.* natural sources for populations in rivers is unknown. Its saprophytic capacity could allow for establishment of “natural” reservoirs but it could also be leaked into rivers from anthropogenic sources. Similarly, for the plant pathogen species complex *Ralstonia solanacearum*, aquatic plants and sediments seem to be reservoirs for populations of this group of bacteria when detected in water (Hong et al., 2008; Tomlinson et al., 2009) suggesting that this group of bacteria, like SRP are probably transients rather than residents in surface waters. These cases beckon the need for further research to find environmental sources to improve the understanding of disease epidemiology. It could be useful to investigate the incorporation of plant pathogenic bacteria into epilithic biofilms, such as been observed for Psy (Morris et al., 2007) where they could be protected by the biofilm matrix and access sugars from algal photosynthesis.

Although it is likely that there is run-off of these two groups of bacteria into the Durance River from vegetation, we observed that water temperature is strongly correlated with the densities of the populations of Psy and SRP in river water: population densities were positively correlated with temperature for SRP and negatively correlated with temperature for Psy. Temperature was correlated with altitude of the site – and this could reflect differences in land use and vegetation type along the banks of the rivers, with production of annual crops typifying the lower basin, fruit orchards in the lower part of the middle basin and forests and unmanaged lands more frequent in the upper basin. Nevertheless, the trend with temperature observed throughout the entire Durance catchment was also observed within each of the three basins (upper, middle and lower) when considered separately. This further strengthens the hypothesis that temperature is a critical factor and not simply a reflection of its correlation with other gradients across the entire catchment. Temperature seems to influence the abundance of these two groups of bacteria whatever the context of the basin and the associated sources of bacteria that the river encounters as it crosses different land uses from pastures, to fruit tree production and to vegetable crops with their varying anthropogenic characteristics. The influence of temperature on Psy and SRP appears to be much stronger than on the total culturable bacterial population. This is probably due to differences among the component species in their sensitivity to environmental factors. When they were detected, Psy populations constituted only 10^−5^ % to less than 4% of the total culturable bacterial population and SRP constituted only 10^−5^ % to less than 0.7% of the total culturable population. Therefore, it is reasonable to assume that the perceptible effect of environmental factors on total culturable populations is strongly influenced by the major species components rather than by Psy and SRP. Indeed, the bacterial assemblages in Durance river water are highly diverse when assessed by 16S community profiling (Pédron et al., 2020). The diversity of total bacteria suggests that there is not only a range of environmental tolerances among the bacteria in the river system that makes it difficult to identify overriding correlations with environmental factors, but that there are also opportunities for competition and antagonistic interactions. However, when there were significant correlations between densities of total culturable bacterial populations and those of Psy or SRP, they were positive suggesting that increasing densities of bacteria that co-occur with Psy or SRP in river water were not detrimental to the populations of these latter two groups of bacteria. Such positive correlations between Psy and total culturable bacteria in water were also observed in a previous study in the Durance catchment (Monteil et al., 2013). Furthermore, a metagenomic analysis of samples from three sites along the Durance catchment representing the upper-, mid- and lower basins (Pédron et al., 2020) showed the same trend for the *Pseudomonas* genus as we observed for the *P. syringae* complex suggesting that *P. syringae* might be representative of the genus as a whole in terms of its population dynamics in river water.

A remarkable observation for both Psy and SRP is that river water contains a diversity of populations of these groups of bacteria beyond what is known to be associated with disease on crops. This raises the intriguing question of the origin of these bacteria in river water. For example, river water harbors genetic groups of Psy and SRP with no known epidemiological importance - PG10 and PG13 for Psy (Berge et al., 2014) and *P. aquaticum* and *P. quasiaquaticum* for SRP (Pédron et al., 2019; Portier et al., 2020; Ben Moussa et al., 2021). Some strains of Psy PG10 and PG13 have been found in association with plants (Borschinger et al., 2015), but the vast part of their diversity has been found in water elsewhere (Berge et al., 2014). The recently discovered *P. aquaticum* and *P. quasiaquaticum* found in the Durance catchment have been reported only from aquatic environments and have never been isolated from plant since 1944 in contrast to other all the other *Pectobacterium* species deposited since 1944 at the CIRM-CFBP bacterial collection (Portier et al., 2020). Perhaps it is autochthonous in water but the low prevalence of SRP in water suggests it has other yet-to-be-discovered habitats that serve as sources for populations in the river. Notably, *P. aquat*icum was found mostly in the lower half of the Durance catchment with an important occurrence on a limited number of sites suggesting its requirement for either very specific conditions or its association with a limited number of sources. This is in contrast to *P. versatile*, the most represented *Pectobacterium* species deposited to the CIRM-CFBP collection since 1944, known to be associated with a wide range of plant species (crops and ornamentals, for example) (Portier et al., 2020), that was detected throughout the catchment.

River water also contained strains that are likely to be of epidemiological importance – but not necessarily in the Durance River catchment or on crops. We detected *P. peruviense* – with the same techniques described in this work - (Ben Moussa et al., 2022), a species that has only been reported at high altitudes in South America as a pathogen of potato (Waleron et al., 2018). Its presence in the Durance River suggests that it has a natural but previously unknown ubiquity in the environment or that there was a rare and unrecognized dissemination event from South America. The presence of a few strains of *P. atrosepticum*, a species mostly recorded on potato (Ben Moussa et al., 2022), also raises the question of its origin – either from disease on the very small surface of potato crops in the Durance catchment or the association of this bacterium with wild solanaceous plants or a few brassicas (Toth et al., 2021). Other plant sources are unlikely for this species in light of its very narrow host range (Ma et al., 2007; Toth et al., 2021). Furthermore, we wonder if the presence in river water of *D. oryzae* (Ben Moussa et al., 2022), known to be a pathogen of rice but also pathogenic on potato, maize and several other crops (Hugouvieux-Cotte-Pattat & Van Gijsegem, 2021), might have its origin in disease on the small amount of regional potato crop or an association with wild grasses that has yet to be described. For Psy, its population is dominated by a ubiquitous subgroup of PG02 (here named *cts*-haplotype DD.1) that has been associated with numerous crop disease epidemics. Nevertheless, the epidemics linked to the DD.1 haplotype of Psy have occurred mostly outside of the Durance River basin with the closest known epidemic in Southwestern France (Morris et al., 2000). For another haplotype in PG02 closely related to DD.1, quasi-clonal lines from epidemics of cantaloupe blight in southwestern France, from snowfall in the French Vercors Massif and from a pristine creek on the south island of New Zealand have been identified (Monteil et al., 2016) supporting the hypothesis that long distance movement of bacteria – even between the northern and southern hemispheres – does occur. Hence, there are indeed mechanisms for long distance movement - most likely via the atmosphere - that can link rivers with cropped fields elsewhere.

To understand the potential epidemiological significance of the presence of diverse Psy and SRP throughout the Durance catchment, we need to identify the processes that have contributed to this state of the microbiology of river water. We wonder if the assemblages of Psy and SRP populations in river water are the result of rivers being simply collectors of bacteria from the local landscape (from run-off, for example) and from more distant sources (via rain and snowfall, for example). If run-off is the main process leading to the abundance and diversity of Psy and SRP in river water, it would be very important to identify all of the potential sources including prairies, pastures and wild plant stands in addition to known crop hosts for disease. It is also important to consider if Psy and SRP simply survive or if there is multiplication and diversification. Interestingly, pathogenicity tests with *D. dianthicola*-like strains isolated from river water in Finland revealed that water-borne strains were more aggressive than strains of *D. dianthicola* isolated from potato (Laurila et al., 2008). Furthermore, *D. aquatica* isolated in Finnish rivers were later found to be aggressive on acidic fruits such as tomato or cucumber (Duprey et al., 2019). The lack of xylanases and xylose degradation pathways in *D. aquatica* could reflect adaptation to aquatic charophyte hosts which, in contrast to land plants, do not contain xyloglucans. This suggests that water-borne species have experienced some selective pressures that lead to adaptations that could, in turn, be useful in causing disease to crops.

Our results point to the need to clarify the role of temperature in influencing population densities of Psy and SRP. The differential effect of temperature on population sizes of Psy and SRP could be due to effects on growth and/or die-off – both processes being important in structuring the gene pool of these populations. In laboratory tests of growth of SRP strains inoculated into filter- and autoclave-sterilized river water, 100-fold increases in populations of *Dickeya* and *Pectobacterium* strains were observed over 10 days at 20°c. However, at 8°C growth was lower for *Pectobacterium* and die-off was observed for *Dickeya* (Ben Moussa et al., 2022). This implies that, under natural conditions, there are stresses caused by a fluctuating environment that maintain SRP populations at low levels. Our work provides insight into what these stresses might be. For Psy, a previous study suggested that populations in rivers did not necessarily multiply (Monteil et al., 2013). The authors of that study noted that the similarities in population structure between rain, snow melt and headwaters in France could be attributed to effective transportation of Psy strains with snow melt and rain water infiltrating through the soil of subalpine grasslands. However, in a study of headwaters in France, the USA and New Zealand it was observed that about half of the populations at the headwater sites were composed of *cts* haplotypes that were unique to the region from which they were sampled (43% for New Zealand headwaters, 67% for USA headwaters and 70% for French headwaters) (Morris et al., 2010), implying the existence of a local diversification process. Preliminary laboratory experiments show that growth in river water is possible (Berge, unpublished) thereby suggesting that this could contribute to diversification.

A critical epidemiological aspect of the regular occurrence of Psy and SRP in the Durance River catchment is the potential of river-borne bacteria to cause disease to crops irrigated with river water. This concern brings to the forefront the questions of how to assess the epidemiological potential of river-borne bacteria and how to anticipate disease outbreaks. The epidemiological potential of Psy and SRP strains in river water could be addressed via pathogenicity tests such as those conducted for the *D. dianthicola*-like and *D. aquatica* strains isolated from river water in Finland (Laurila et al., 2008). However, in the case of Psy, the choice of pertinent hosts to test against strains in the dominant DD.1 haplotype is complicated by its variable and potentially broad host range (Morris et al., 2019). To anticipate disease outbreaks, data on epidemiological potential needs to be set in the context of rate of exposure of crop plants and the local environmental conditions. Exposure of plants due to irrigation with river water could be estimated. For the main departments of France that irrigate with water from the Durance River catchment (Alpes de Haute Provence, Hautes Alpes and Vaucluse) there are > 55000 ha of agriculture that could be irrigated including fruits and vegetables, pastures and cereals (Chambre_d’Agriculture_PACA, 2014). In the case of lettuce – a vegetable crop produced in abundance in the Durance River catchment - plants require a total of about 30 mm (30 L m^−2^) in the few days following planting (about a week) and about 5 or 6 subsequent irrigations of about 15 mm each during the 2 to 3 months of culture afterwards for a total of about 110 mm (110 L m^−2^) during the life cycle of the plant (Lecompte, 2012). For a lettuce field planted at a density of 150000 plants ha^−1^ and a mean bacterial population density of 10^3^ Psy or SRP L^−1^ of river water, each plant on average could potentially be in contact with more than 10^3^ bacteria belonging to the Psy or SRP species complex from water during the first few days after planting and about 10^4^ Psy or SRP bacteria throughout the period of culture due to irrigation. At a first glance, this might seem to be cause for alarm. However, the fate of these bacteria is unknown. We do not know if they survive, if they are physiologically competent, or if they are compatible with the crop they encounter. We should keep in mind that only a fraction of the Psy or SRP strains – or perhaps none at all - that contact the plants via irrigation will have pathogenic potential for the crop that they encounter. Overall, this speculation points out why it is important to have quantitative data on bacterial population size that allows for estimations of exposure - and that go beyond the uncertainties of risk assessment based simply on presence and prevalence.

Historical epidemiological information for the Durance River catchment does not point to Psy and SRP as re-occurring pathogens of crops in this region - with the exception of bacterial canker of apricot (Parisi et al., 2019) and leaf spot of lettuce (Allex & Rat, 1990), both caused by Psy. This could suggest that the range of environmental conditions, the historical land use and the intensity of agriculture up to present are within the spectrum of conditions that do not generally favor epidemics by Psy or SRP. It could also suggest that the bacteria present in the two species complexes studied here are not well adapted to the cultivated crops cultivated in the Durance River catchment. In light of these observations, we can make recommendations that could contribute to creating indicators of the risk posed by river-borne populations of Psy and SRP. These indicators could combine various variables that account for i) total population sizes of Psy and SRP (or the dominant genetic lines), ii) water temperature and iii) the various chemical conditions in each of the three basins that we determined to be correlated with Psy and SRP population densities. Risk alerts could be developed to express the deviation from the trends we observed here. These indicators could be confronted with previsions for major changes in agricultural land use such as changes in the geographic ranges of certain crops or the introduction of new crops into the region. This would require assessment of their sensitivity to diseases caused by Psy and SRP in the environmental conditions associated with the anticipated changes.

This work raises the general question of how river water reflects the diversity of plant-associated microorganisms beyond what is reflected by populations associated with crops or other vegetation. We raise the critical question of how this diversity be used for anticipating disease emergence and the need to elucidate the underlying processes that connect these populations to epidemics. These processes would indeed be targets for management. Integrating nonagricultural reservoirs of plant pathogens – such as river water – into a more comprehensive vision of pathogen ecology and life history could improve forecasting disease risk and anticipating epidemics in the face of changes in land use and climate. Although various bacteria and fungi have been detected in irrigation water (Zappia et al., 2014; Lamichhane & Bartoli, 2015), some of these might be present only when inoculum reservoirs in diseased plants are nearby and they might not be able to persist in river water. To develop such an integrative approach, it will be important to distinguish pathogens with the capacity to thrive in environmental reservoirs vs. those whose presence in the environment represents transient residues from agriculture. This approach would open new directions in disease surveillance that would allow for anticipation on larger scales of space and time and could foster better adaptation of land use in the face of changing climate.

## Supporting information

Supplemental_Tabs

## Acknowledgments

This work was possible due to funding contract ANR-17-CE32-0004 for the project on *Strategic preemptive pathogen surveillance of air and water to anticipate plant disease emergence in scenarios of changing land use.* We thank Karine Berthier (INRAE-PACA-Pathologie Végétale) for valuable insights on bioinformatics analyses. We thank Ariane Toussaint for her interest in the project and her logistic support in Val-des-Prés during the sampling campaigns in the upper Durance. We thank the GeT-PlaGe core facility (INRAE, Toulouse, France) for their collaboration, and especially Olivier Bouchez, Arien Castinel, Marie Gislard and Lisa Gil.

## Data, script and code availability

Data are available in the six tables and two figures of Supplemental Information section of this work. Scripts for calculating population densities of the *Pseudomonas syringae* species complex in the Durance river in 2016-2017 are available at https://entrepot.recherche.data.gouv.fr/dataset.xhtml?persistentId=doi:10.57745/TB60SC.

## Supplementary Information

Available at: https://www.biorxiv.org/content/10.1101/2022.09.06.506731v1.supplementary-material

**Supplementary Table 1:** Population sizes of bacteria in the Durance river, tributaries and canals

**Supplementary Table 2:** Description of amplicon sequence variants.

**Supplementary Table 3:** Variability of population sizes of bacteria at two sampling site along the Durance river.

**Supplementary Table 4:** Primers used for NGS for *P. syringae*

**Supplementary Table 5:** *Cts* sequences of reference strains of *Pseudomonas syringae* used in this study

**Supplementary Table 6:** Values for water physical-chemistry variables.

**Supplementary Figure 1**: Relationship between population densities of *Pseudomonas syringae* and Soft Rot Pectobacteriaceae (SRP) species complexes in Durance River

**Supplementary Figure 2**: Variability in densities of *Pseudomonas syringae* and Soft Rot Pectobacteriaceae (SRP) at three sampling times within the same day at two sites along the Durance River catchment.

## References

1. Allex D, Rat B (1990) Les bactérioses des salades: un problème omniprésent. PHM-Rev. Hort., 310, 45–50.

2. Andrew JT, Sauquet E (2017) Climate change impacts and water management adaptation in two Mediterranean-climate watersheds: Learning from the Durance and Sacramento Rivers. Water, 9, doi: 10.3390/w9020126. https://www.mdpi.com/2073-4441/9/2/126

3. Ben Moussa H, Bertrand C, Rochelle-Newall E, Fiorini S, Pédron J, Barny MA (2022) The diversity of soft rot Pectobacteriaceae along the Durance River stream in the south-east of France revealed by multiple seasonal surveys. Phytopathology, 112, 1676–1685. 10.1094/PHYTO-12-21-0515-R

4. Ben Moussa H, Pédron J, Bertrand C, Hecquet A, Barny M-A (2021) *Pectobacterium quasiaquaticum* sp. nov., isolated from waterways. International Journal of Systematic and Evolutionary Microbiology, 71. https://www.microbiologyresearch.org/content/journal/ijsem/10.1099/ijsem.0.005042

5. Berge O, Monteil CL, Bartoli C, Chandeysson C, Guilbaud C, Sands DC, Morris CE (2014) A user’s guide to a data base of the diversity of *Pseudomonas syringae* and its application to classifying strains in this phylogenetic complex. Plos one, 9, (9): e105547. doi:10.5510.101371/journal.pone.0105547. 10.1371/journal.pone.0105547

6. Borschinger B, Bartoli C, Chandeysson C, Guilbaud C, Parisi L, Bourgeay JF, Buisson E, Morris CE (2015) A set of PCRs for rapid identification and characterization of *Pseudomonas syringae* phylogroups. Journal of Applied Microbiology, 120, 714–723. 10.1111/jam.13017

7. Callahan BJ, McMurdie PJ, Rosen MJ, Han AW, Johnson AJA, Holmes SP (2016) DADA2: High-resolution sample inference from Illumina amplicon data. Nature Methods, 13, 581–583. 10.1038/nmeth.3869

8. Chambre_d’Agriculture_PACA. (2014). SRHA Provence Alpes Côte d’Azur: Stratégie Régionale Hydraulique Agricole. https://paca.chambres-agriculture.fr/fileadmin/user_upload/National/FAL_commun/publications/Provence-Alpes-Cote_d_Azur/rdiagnostic-agriculture_irriguee_paca_2014.pdf (accessed 1 April 2022)

9. Cigna J, Dewaegeneire P, Beury A, Gobert V, Faure D (2017) A *gapA* PCR-sequencing assay for identifying the *Dickeya* and *Pectobacterium* potato pathogens. Plant Disease, 101, 1278–1282. 10.1094/PDIS-12-16-1810-RE

10. Duprey A, Taib N, Leonard S, Garin T, Flandrois J-P, Nasser W, Brochier-Armanet C, Reverchon S (2019) The phytopathogenic nature of *Dickeya aquatica* 174/2 and the dynamic early evolution of *Dickeya* pathogenicity. Environmental Microbiology, 21, 2809–2835. https://sfamjournals.onlinelibrary.wiley.com/doi/abs/10.1111/1462-2920.14627

11. Eayre CG, Bartz JA, Concelmo DE (1995) Bacteriophages of *Erwinia carotovora* and *Erwinia ananas* isolated from freshwater lakes. Plant Disease, 79, 801–804

12. Escudié F, Auer L, Bernard M, Mariadassou M, Cauquil L, Vidal K, Maman S, Hernandez-Raquet G, Combes S, Pascal G (2018) FROGS: Find, Rapidly, OTUs with Galaxy Solution. Bioinformatics, 34, 1287–1294. 10.1093/bioinformatics/btx791

13. Faye P, Bertrand C, Pédron J, Barny M-A (2018) Draft genomes of “*Pectobacterium peruviense*” strains isolated from fresh water in France. Standards in Genomic Sciences, 13, 27. 10.1186/s40793-018-0332-0

14. Gardan L, Cottin S, Bollet C, Hunault G (1991) Phenotypic heterogeneity of *Pseudomonas syringae* van Hall. Res. Microbiol., 142, 995–1003. 10.1016/0923-2508(91)90010-8

15. Gorski L, Cooley MB, Oryang D, Carychao D, Nguyen K, Luo Y, Weinstein L, Brown E, Allard M, Mandrell RE, Chen Y (2022) Prevalence and clonal diversity of over 1,200 *Listeria monocytogenes* isolates collected from public access waters near produce production areas on the Central California Coast during 2011 to 2016. Appl Environ Microbiol, 88, e0035722. 10.1128/aem.00357-22

16. Guillorit-Rondeau C, Malandrin L, Samson R (1996) Identification of two serological flagellar types (H1 and H2) in *Pseudomonas syringae* pathovars. EuropeanJournal of Plant Pathology, 102, 99–110. https://link.springer.com/article/10.1007/BF01877120

17. Harrison M, Franc G, Maddox D, Michaud J, Mccarter-Zorner N (1987) Presence of *Erwinia carotovora* in surface water in North America. Journal of Applied Bacteriology 62, 565–570. 10.1111/j.1365-2672.1987.tb02690.x

18. Hélias V, Hamon P, Huchet E, van der Wolf J, Andrivon D (2012) Two new effective semiselective crystal violet pectate media for isolation of *Pectobacterium* and *Dickeya*. Plant Pathology, 61, 339–345. 10.1111/j.1365-3059.2011.02508.x

19. Hong JC, Momol MT, Jones JB, Ji P, Olson SM, Allen C, Perez A, Pradhanang P, Guven K (2008) Detection of *Ralstonia solanacearum* in irrigation ponds and aquatic weeds associated with the ponds in north Florida. Plant Dis, 92, 1674–1682. 10.1094/PDIS-92-12-1674

20. Hugouvieux-Cotte-Pattat N, Van Gijsegem F (2021) Diversity within the *Dickeya zeae* complex, identification of *Dickeya zeae* and *Dickeya oryzae* members, proposal of the novel species *Dickeya parazeae* sp. nov. International Journal of Systematic and Evolutionary Microbiology, 71. 10.1099/ijsem.0.005059

21. Ivanović Ž, Blagojević J, Nikolić I (2018) Leaf spot disease on *Philodendron scandens, Ficus carica* and *Actinidia deliciosa* caused by *Pseudomonas syringae* pv. *syringae* in Serbia. European Journal of Plant Pathology, 151, 1107–1113. 10.1007/s10658-018-1437-4

22. James E, Joyce M (2004) Assessment and management of watershed microbial contaminants. Critical Reviews in Environmental Science and Technology, 34, 109–139. 10.1080/10643380490430663

23. Jorge PE, Harrison MD (1986) The association of *Erwinia carotovora* with surface water in northeastern Colorado. I. The presence and population of the bacterium in relation to location, season and water temperature. American Potato Journal, 63, 517–531. https://link.springer.com/article/10.1007/BF03044052

24. King EO, Ward MK, Raney DE (1954) Two simple media for the demonstration of pyocyanin and fluorescein. J. Lab. & Clin. Med., 44, 301–307

25. Kuentz A (2013) Un siècle de variabilité hydro-climatique sur le bassin de la Durance: Recherches historiques et reconstitutions. AgroParisTech, Paris. https://pastel.archives-ouvertes.fr/tel-01171004/document

26. Lamichhane JR, Bartoli C (2015) Plant pathogenic bacteria in open irrigation systems: what risk for crop health? Plant Pathology, 64, 757–766. 10.1111/ppa.12371

27. Laurila J, Ahola V, Lehtinen A, Joutsjoki T, Hannukkala A, Rahkonen A, Pirhonen M (2008) Characterization of *Dickeya* strains isolated from potato and river water samples in Finland. European Journal of Plant Pathology, 122, 213–225. 10.1007/s10658-008-9274-5

28. Lecompte F (2012) Management of soil nitrate heterogeneity resulting from crop rows in a lettuce–tomato rotation under a greenhouse. Agronomy for Sustainable Development, 32, 811–819. 10.1007/s13593-011-0047-8

29. Lindeberg M, Cunnac S, Collmer A (2012) *Pseudomonas syringae* type III effector repertoires: last words in endless arguments. Trends in Microbiology, 20, 199–208. 10.1016/j.tim.2012.01.003

30. Ma B, Hibbing ME, Kim HS, Reedy RM, Yedidia I, Breuer J, Breuer J, Glasner JD, Perna NT, Kelman A, Charkowski AO (2007) Host range and molecular phylogenies of the soft rot enterobacterial genera *Pectobacterium* and *Dickeya*. Phytopathology, 97, 1150–1163. 10.1094/PHYTO-97-9-1150

31. McCarter-Zorner NJ, Franc GD, Harrison MD, Michaud JE, Quinn CE (1984) Soft rot Erwinia bacteria in surface and underground waters in southen Scotland and in Colorado, United-States. J. Appl. Bacteriol., 57, 95–105. 10.1111/j.1365-2672.1984.tb02361.x

32. Monteil CL, Bardin M, Morris CE (2014) Features of air masses associated with the deposition of Pseudomonas syringae and Botrytis cinerea by rain and snowfall. ISME J, 8, 2290–2304. https://www.nature.com/articles/ismej201455

33. Monteil CL, Lafolie F, Laurent J, Clement J-C, Simler R, Travi Y, Morris CE (2013) Soil water flow is a source of the plant pathogen *Pseudomonas syringae* in subalpine headwaters. Environ. Microbiol., 16, 2038– 2052. 10.1111/1462-2920.12296

34. Monteil CL, Yahara K, Studholme DJ, Mageiros L, Méric G, Swingle B, Morris CE, Vinatzer BA, Sheppard SK (2016) Population-genomic insights into emergence, crop-adaptation, and dissemination of *Pseudomonas syringae* pathogens. Microbial Genomics, doi: 10.1099/mgen.0.000089. 10.1099/mgen.0.000089

35. Morgan M, Anders S, Lawrence M, Aboyoun P, Pagès H, Gentleman R (2009) ShortRead: a bioconductor package for input, quality assessment and exploration of high-throughput sequence data. Bioinformatics, 25, 2607–2608. 10.1093/bioinformatics/btp450

36. Morris CE, Glaux C, Latour X, Gardan L, Samson R, Pitrat M (2000) The relationship of host range, physiology, and genotype to virulence on cantaloupe in *Pseudomonas syringae* from cantaloupe blight epidemics in France. Phytopathology, 90, 636–646. 10.1094/PHYTO.2000.90.6.636

37. Morris CE, Kinkel LL, Kun X, Prior P, Sands DC (2007) A surprising niche for the plant pathogen *Pseudomonas syringae*. Infection, Genetics and Evolution, 7, 84–92. 10.1016/j.meegid.2006.05.002

38. Morris CE, Lamichhane JR, Nikolić I, Stanković S, Moury B (2019) The overlapping continuum of host range among strains in the *Pseudomonas syringae* complex. Phytopathology Research, 1, 4. 10.1186/s42483-018-0010-6

39. Morris CE, Monteil CL, Berge O (2013) The life history of *Pseudomonas syringae*: linking agriculture to Earth system processes. Annu. Rev. Phytopathol., 51, 85–104. 10.1146/annurev-phyto-082712-102402

40. Morris CE, Sands DC, Vanneste JL, Montarry J, Oakley B, Guilbaud C, Glaux C (2010) Inferring the evolutionary history of the plant pathogen *Pseudomonas syringae* from its biogeography in headwaters of rivers in North America, Europe and New Zealand. mBio, 1(3): e00107-10-e00107-20. 10.1128/mBio.00107-10

41. Parisi L, Morgaint B, Blanco-Garcia J, Guilbaud C, Chandeysson C, Bourgeay JF, Moronvalle A, Brun L, Brachet ML, Morris CE (2019) Bacteria from four phylogroups of the *Pseudomonas syringae* complex can cause bacterial canker of apricot. Plant Pathology, 68, 1249–1258. 10.1111/ppa.13051

42. Parkinson N, Bryant R, Bew J, Conyers C, Stones R, Alcock M, Elphinstone J (2013) Application of variable-number tandem-repeat typing to discriminate *Ralstonia solanacearum* strains associated with English watercourses and disease outbreaks. Applied and environmental microbiology, 79, 6016–6022. 10.1128/AEM.01219-13

43. Pédron J, Bertrand C, Taghouti G, Portier P, Barny M-A (2019) *Pectobacterium aquaticum* sp. nov., isolated from waterways. International Journal of Systematic and Evolutionary Microbiology, 69, 745–751. 10.1099/ijsem.0.003229

44. Pédron J, Guyon L, Lecomte A, Blottière L, Chandeysson C, Rochelle-Newall E, Raynaud X, Berge O, Barny M-A (2020) Comparison of environmental and culture-derived bacterial communities through 16S metabarcoding: A powerful tool to assess media selectivity and detect rare taxa. Microorganisms, 8, 1129. https://www.mdpi.com/2076-2607/8/8/1129

45. Pérombelon MCM, Kelman A (1980) Ecology of the soft rot Erwinias. Ann. Rev. Phytopathol., 18, 361–387. 10.1146/annurev.py.18.090180.002045

46. Pietsch RB, Vinatzer BA, Schmale DG (2017) Diversity and abundance of ice nucleating strains of *Pseudomonas syringae* in a freshwater lake in Virginia, USA. Frontiers in Microbiology, 8. 10.3389/fmicb.2017.00318

47. Portier P, Pédron J, Taghouti G, Dutrieux C, Barny M-A (2020) Updated taxonomy of *Pectobacterium* genus in the CIRM-CFBP bacterial collection: When newly described species reveal “old” endemic population. Microorganisms, 8, 1441. 10.3390/microorganisms8091441

48. Potrykus M, Golanowska M, Sledz W, Zoledowska S, Motyka A, Kolodziejska A, Butrymowicz J, Lojkowska E (2015) Biodiversity of *Dickeya* spp. isolated from potato plants and water sources in temperate climate. Plant Disease, 100, 408–417. 10.1094/PDIS-04-15-0439-RE

49. R_Core_Team (2020) R: A language and environment for statistical computing. R Foundation for Statistical Computing, Vienna, Austria., https://www.R-project.org/.

50. Rankinen K, Butterfield D, Faneca Sànchez M, Grizzetti B, Whitehead P, Pitkänen T, Uusi-Kämppä J, Leckie H (2016) The INCA-Pathogens model: An application to the Loimijoki River basin in Finland. Science of The Total Environment, 572, 1611–1621. 10.1016/j.scitotenv.2016.05.043

51. Sarkar SF, Guttman DS (2004) Evolution of the core genome of *Pseudomonas syringae*, a highly clonal, endemic plant pathogen. Appl. Environ. Microbiol., 70, 1999–2012. 10.1128/AEM.70.4.1999-2012.2004

52. Stopelli E, Conen F, Guilbaud C, Zopfi J, Alewell C, Morris CE (2017) Ice nucleators, bacterial cells and *Pseudomonas syringae* in precipitation at Jungfraujoch. Biogeosciences, 14, 1189–1196. 10.5194/bg-14-1189-2017

53. Tomlinson DL, Elphinstone JG, Soliman MY, Hanafy MS, Shoala TM, Abd El-Fatah H, Agag SH, Kamal M, Abd El-Aliem MM, Fawzi FG, Stead DE, Janse JD (2009) Recovery of *Ralstonia solanacearum* from canal water in traditional potato-growing areas of Egypt but not from designated Pest-Free Areas (PFAs). European Journal of Plant Pathology, 125, 589–601. 10.1007/s10658-009-9508-1

54. Toth IK, Barny M-a, Brurberg MB, Condemine G, Czajkowski R, Elphinstone JG, Helias V, Johnson SB, Moleleki LN, Pirhonen M, Rossmann S, Tsror L, van der Waals JE, van der Wolf JM, Van Gijsegem F, Yedidia I (2021) *Pectobacterium* and *Dickeya*: Environment to Disease Development. In: Plant Diseases Caused by Dickeya and Pectobacterium Species eds Van Gijsegem F, van der Wolf JM, & Toth IK), pp. 39–84. Springer International Publishing, Cham. 10.1007/978-3-030-61459-1_3

55. Waleron M, Misztak A, Waleron M, Franczuk M, Wielgomas B, Waleron K (2018) Transfer of *Pectobacterium carotovorum* subsp. *carotovorum* strains isolated from potatoes grown at high altitudes to *Pectobacterium peruviense* sp. nov. Systematic and Applied Microbiology, 41, 85–93. 10.1016/j.syapm.2017.11.005

56. Whitehead PG, Leckie H, Rankinen K, Butterfield D, Futter MN, Bussi G (2016) An INCA model for pathogens in rivers and catchments: Model structure, sensitivity analysis and application to the River Thames catchment, UK. Science of The Total Environment, 572, 1601–1610. 10.1016/j.scitotenv.2016.01.128

57. Wickham H (2016) ggplot2: Elegant Graphics for Data Analysis. Springer-Verlag, New York. https://ggplot2.tidyverse.org

58. Zappia RE, Hüberli D, Hardy GESJ, Bayliss KL (2014) Fungi and oomycetes in open irrigation systems: knowledge gaps and biosecurity implications. Plant Pathology, 63, 961–972. 10.1111/ppa.12223

