## Supplemental_Figs for "Comparative abundance and diversity of populations of the *Pseudomonas syringae* and Soft Rot *Pectobacteriaceae* species complexes throughout the Durance River catchment from its French Alps sources to its delta": Morris_Durance_SuppFig_01.docx

**Supplementary Figure 1**: Relationship between population densities of *Pseudomonas syringae* and Soft Rot Pectobacteriaceae (SRP) species complexes in Durance River water for 87 samples collected in 2016 and 2017 in which the two groups of bacteria occurred together in the same sample. The dotted line indicates equivalent population densities for the two groups of bacteria.

*Morris et al. 2022. P. syringae* and Soft Rot *Pectobacteriaceae* in the Durance River catchment

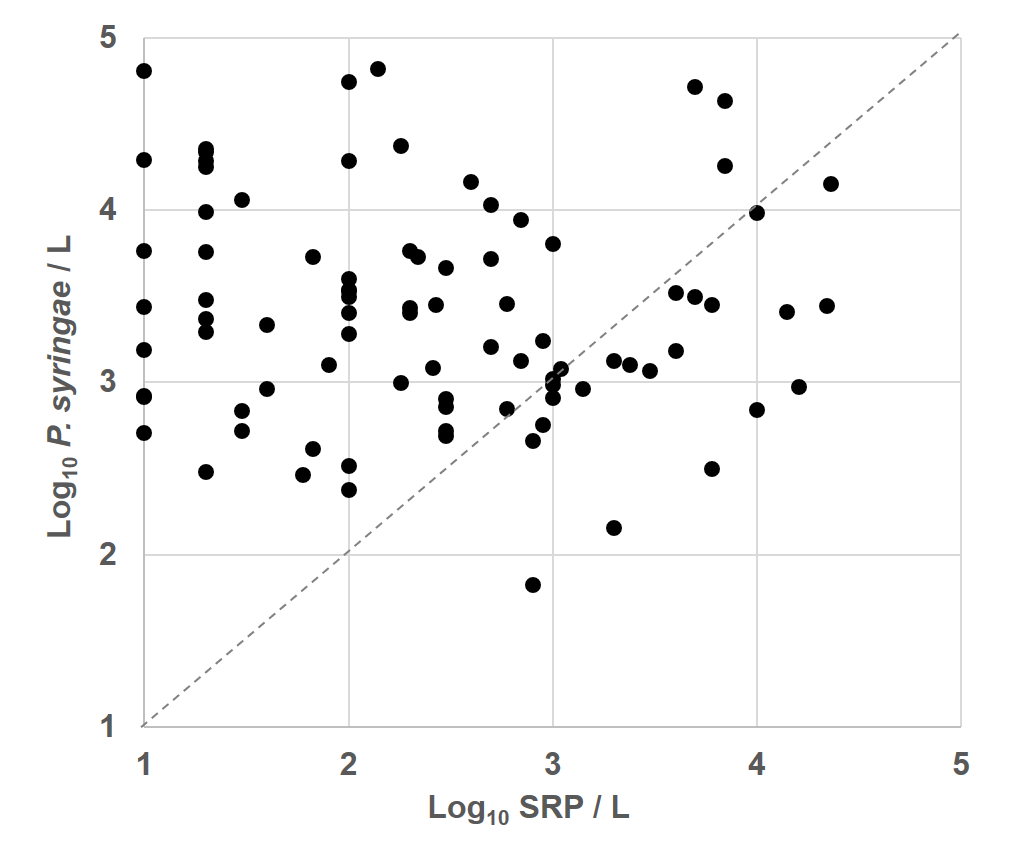
