## Supplemental_Figs for "Comparative abundance and diversity of populations of the *Pseudomonas syringae* and Soft Rot *Pectobacteriaceae* species complexes throughout the Durance River catchment from its French Alps sources to its delta": Morris_Durance_SuppFig_02.docx

**Supplementary Figure 2**: Variability in densities of *Pseudomonas syringae* and Soft Rot Pectobacteriaceae (SRP) at three sampling times within the same day at two sites along the Durance River catchment.

Morris et al. 2022. *P. syringae* and Soft Rot Pectobacteriaceae in the Durance River catchment

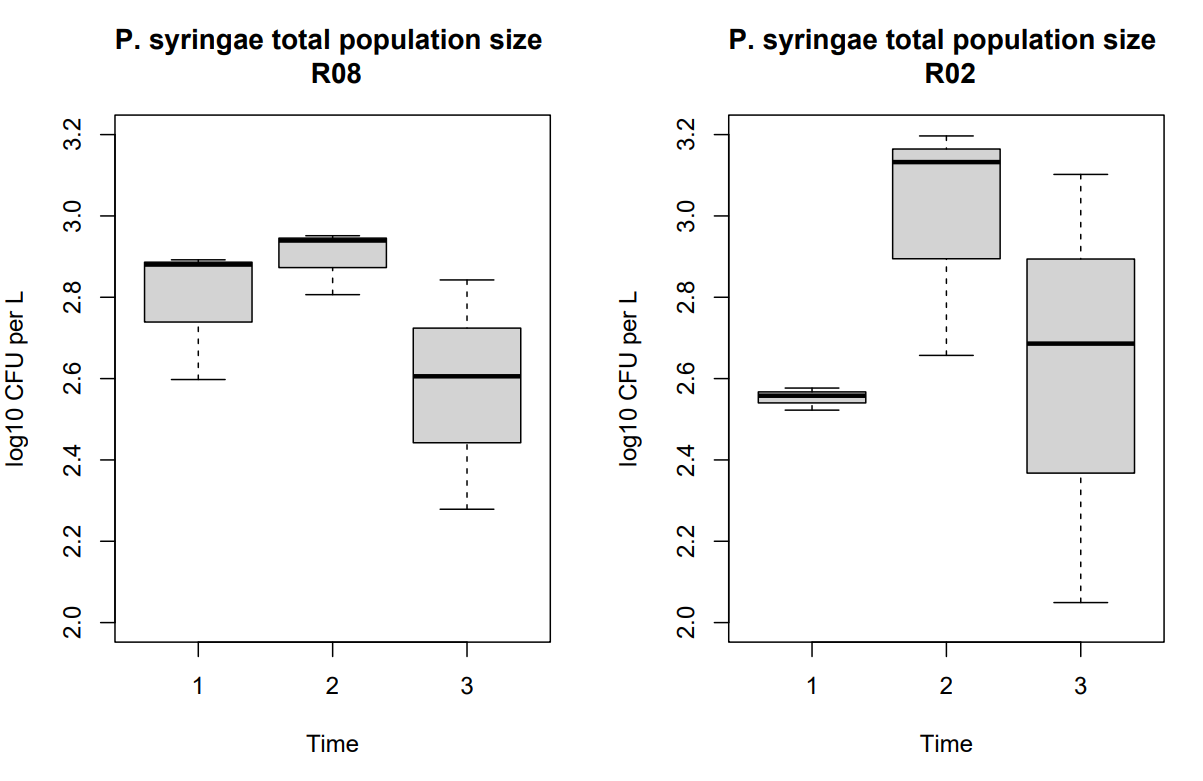

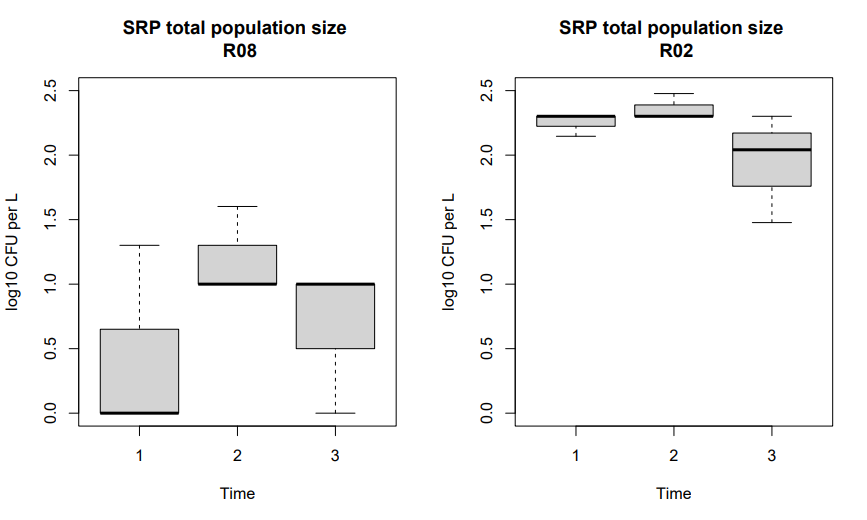
